# Interplay of protein disorder in retinoic acid receptor heterodimer and its corepressor regulates gene expression

**DOI:** 10.1101/452482

**Authors:** Tiago N. Cordeiro, Nathalie Sibille, Pierre Germain, Philippe Barthe, Abdel Boulahtouf, Fréderic Allemand, Rémy Bailly, Valérie Vivat, Christine Ebel, Alessandro Barducci, William Bourguet, Albane le Maire, Pau Bernadó

## Abstract

The retinoic acid receptors (RARs) form heterodimers with retinoid X receptors (RXRs) and control gene transcription in response to ligand binding and *via* allosteric activation of the C-termini helix (helix H12) of its ligand-binding domain. Herein we show that in the absence of ligand, helices H12 of RXR and RAR are disordered. The selective RAR agonist, Am580, induces folding of H12, whereas in the presence of the inverse agonist BMS493, H12 stays mostly disordered. These results substantiate a link between the structural dynamics of H12 and RXR/RAR heterodimer biological functions, and highlight disordered-to-order transition as an essential mechanism for retinoic acid mediated regulation. Unliganded RAR exerts a strong repressive activity allowed by the recruitment of transcriptional corepressors and establishment of a corepressor complex in the promoter region of target genes. The human regulatory complex of the RARα bound to the full-length interaction domain of the corepressor N-CoR was studied by integrating several experimental (SAXS, X-ray crystallography, NMR, CD, AUC) and computational data. Unexpectedly, we found that, while mainly intrinsically disordered, the N-CoR presents partially evolutionary conserved structured regions that are involved in transient intramolecular contacts. In the presence of RXR/RAR, we show that N-CoR exploits its multivalency to form a multi-site complex that diplays an equilibrium between different conformational states. This conformational equilibrium is modulated by cognate ligands, RAR point mutation and RXR H12 deletion. Now, we can state that, in addition to NR conformation and ligand-induced allosteric changes, intrinsic disorder is substantially embedded in the synergetic regulation of RXR/RAR activity and its resulting abilities to communicate with the intracellular components.

## Introduction

Nuclear Receptors (NRs) are transcription factors that have a direct role in regulating the expression of ligand-responsive genes. This regulatory capacity of NRs occurs through their ability to recognize specific sequences in the promoters of their target genes and their relationships with the RNA polymerase II holocomplex as well as the chromatin environment that surrounds these genes (Roeder, 1998). Like many other members of the NR family, retinoic acid receptors (NR1B1 (RARα), NR1B2 (RARβ) and NR1B3 (RARγ)) form heterodimeric complexes with the retinoid X receptors (NR2B1 (RXRα), NR2B2 (RXRβ) and NR2B3 (RXRγ)) and function as ligand (retinoic acid)-regulated transcription factors (Gronemeyer et al., 2004; Perissi and Rosenfeld, 2005). In fact, RXR/RAR heterodimers may act either as repressors or activators of gene transcription depending on their ligation status that in turn determines the ability of these DNA-bound receptors to recruit coregulators (either corepressors or coactivators) to target gene promoters (Perissi and Rosenfeld, 2005). Coactivator recruitment is usually ligand-dependent, whereas corepressors interact in most cases with unliganded (apo) receptors.

RARs and RXRs have a conserved modular structure with an N-terminal activation function (AF-1), a central DNA-binding domain (DBD) and a C-terminal ligand-binding domain (LBD) (Germain et al., 2006). The multifunctional LBD is responsible for ligand binding and dimerization and contains a ligand-dependent activation function (AF-2), which corresponds to coregulator interaction surfaces that can be modulated by natural (e.g. retinoic acid) or pharmacological ligands (Gronemeyer et al., 2004). Over the years, different classes of synthetic ligands have been generated to induce or repress gene transcription. While agonists enhance the recruitment of coactivators and destabilize interaction with corepressors, thus inducing the transcription of target genes, inverse agonists do the opposite, decreasing the basal transcriptional activity of apo-receptors (Germain et al., 2002; le Maire et al., 2012). Neutral antagonists inhibit both interactions and block the receptor in an inactive conformation.

In the absence of ligand or in the presence of inverse agonists, RARα (named RAR hereafter) exhibits strong repressive activity that is brought about by the recruitment of corepressors (Glass and Rosenfeld, 2000; McKenna et al., 1999). The two main corepressors, Nuclear receptor CoRepressor (N-CoR/NCoR1/RIP13) (Hörlein et al., 1995) and the Silencing Mediator of Retinoic acid receptor and Thyroid hormone receptors (SMRT/NCoR2/TRAC) (Chen and Evans, 1995), have been shown to reside in, or recruit, high molecular weight complexes that display histone deacetylase activity (Heinzel et al., 1997; Nagy et al., 1997). Deacetylated histones are associated with silent regions of the genome, and it is generally accepted that histone acetylation and deacetylation shuffle nucleosomal targets between a relaxed and condensed chromatin configuration, the former being requisite for transcriptional activation. From a mechanistic point of view, ligand binding to receptors induces a rearrangement of the C-terminal region of the LBD, the so-called helix H12, leading to corepressor dissociation and coactivator recruitment. Coactivators, such as those of the TIF-2/SRC-1/RAC3 (p160) family, mediate the interaction of coactivator complexes with NRs. CBP, p300, P/CAF, and some p160 coactivators themselves are reported to act as histone acetyltransferases (HATs) (Glass and Rosenfeld, 2000; Lonard and O’Malley, 2007). They are capable of acetylating specific residues in the N-terminal tails of different histones, a process that is believed to play an important role in the opening of chromatin during transcription activation (Chen and Evans, 1995; Imhof et al., 1997).

Like many proteins in signaling pathways, coregulators are mainly disordered proteins that act as a platform where multiple proteins attach to perform activities linked to gene transcription (Csizmok et al., 2016; Hegyi et al., 2007). These interactions are mediated by Short Linear Motifs (SLiMs) that are embedded in the sequence and that enable the simultaneous recognition of multiple partners (Van Roey et al., 2014). Regarding the interaction with NRs, two major conserved corepressor/NR recognition motifs (CoRNR box 1-2) LxxI/HIxxxI/L have been identified in SMRT and N-CoR (Hu and Lazar, 1999; Nagy et al., 1999; Perissi et al., 1999). The boxes, which are close in corepressor sequences, define the NR interaction domain (NID).

Important information regarding the molecular mechanisms that regulate the alternative interactions of RAR LBD, with either class of cofactors has been decoded by crystallographic studies using short coregulator-derived peptides (Bourguet et al., 2000; Le Maire et al., 2010; Pogenberg et al., 2005). However, multi-protein complexes containing intrinsically disordered segments, such as the NID of N-CoR, have an extraordinary structural heterogeneity, which poses significant technical challenges for their structural characterization. Indeed, X-ray crystallography is best suited for studies of rigid folded domains and tightly bound complexes. The application of solution Nuclear Magnetic Resonance (NMR), which has been extensively used to study disordered proteins (Dyson and Wright, 2004; Jensen et al., 2013), encounters severe limitations when characterizing large biomolecular complexes. Lower resolution methods can synergistically complement these high-resolution techniques. In particular, small angle X-ray scattering (SAXS) offers a source of structural and dynamic information for highly flexible biomolecules (Bernadó and Svergun, 2012; Cordeiro et al., 2017a). Hybrid approaches, which integrate information from these different techniques into computational tools are the most promising strategy for the structural characterization of highly dynamic proteins and complexes in solution. Here, we report on the structural and dynamic details of the RXR/RAR heterodimer, the disordered NCoR NID, and the highly flexible complex that they form, by integrating solution techniques and computational methods.

Our study reveals that, in addition of the two CoRNR motifs, a conserved region of the NID is partially structured and forms transient intramolecular contacts. Furthermore, we show that NID binds to RXR/RAR through both CoRNR motifs in a highly cooperative manner inducing an equilibrium between several conformational states. Perturbations on the individual CoRNR/NR affinities using RAR ligands and point mutations have a global effect on the structure, dynamics and thermodynamics of the complex. Moreover, we demonstrate that although both receptors contribute to the interaction with N-CoR NID, RAR plays a dominant role over RXR. Thus, we report novel insights into the structural basis of the recruitment of corepressors by RXR/RAR heterodimer, emphasizing the interplay of N-CoR and H12 helix disorder-to-order transitions in the regulation of NR-mediated gene transcription.

## Results

### Disorder in RXR/RAR H12 helices is modulated by ligands and mutations

The most remarkable observation of the multiple crystallographic structures of RXR and RAR reported so far is the conformational variability of their C-terminal helices H12 depending on the ligands and cofactors bound (Bourguet et al., 1995, 2000; Chandra et al., 2017; Germain et al., 2002, 2009; Le Maire et al., 2010; Pogenberg et al., 2005; Renaud et al., 1995; Sato et al., 2010). Nuclear Magnetic Resonance (NMR) and fluorescence anisotropy have also shown the plasticity of these helices in solution depending on the ligation state (Lu et al., 2006; Nahoum et al., 2007). Here, we have used SAXS to validate these previous observations and to analyze other non-crystallized conditions.

SAXS data indicate that unliganded RXR/RAR heterodimer is a globular particle in solution with a radius of gyration, *R_g_* of 26.6 ± 0.4 Å and a maximum intramolecular distance, *D_max_* of 89.0 ± 3.0 Å (Table S1; Fig. 1A). Molecular weight estimation suggests that the particle is a heterodimer, in line with Sedimentation Velocity Analytical Ultracentrifugation (SV-AUC) experiments, *s*_20w_ = 3.78 ± 0.13 S and *f*/*f*_min_= 1.30 ± 0.05 (Fig. S2). The smooth asymmetrical pair-wise distance distribution, *P(r)*, suggests the presence of moderate flexibility in the RXR/RAR heterodimer (Fig. 1A). Crystallographic structures of RXR/RAR (PDB entries 1DKF (Bourguet et al., 2000), 1XDK (Pogenberg et al., 2005), and 3A9E (Sato et al., 2010) (Table S2)), all obtained in the presence of agonist or antagonist of RXR and RAR, were not consistent with measured SAXS data (χ_i_^2^ = 2.55, 2.34 and 1.83, respectively). We hypothesized that the observed discrepancy can be attributed to two reasons: firstly, the missing flexible regions, including the N- and C-termini and the loops connecting helices 2 and 3 in both RXR and RAR (Fig. S1B), and secondly, the positions of RXR and RAR H12 helices that are in antagonist or agonist-bound conformations in the X-ray structures. To validate this hypothesis and based on several crystallographic structures, we built three ensemble models for the RXR/RAR heterodimer in which both H12 helices were maintained disordered, in agonist or in antagonist conformation (see Fig. S1 and methods section for details). SAXS curves derived from these ensemble models were compared with the experimental one. As observed in Figure 1 and Figure S1, the average scattering profile computed from the ensemble model with disordered H12 fragments yielded an excellent agreement to the experimental curve (χ^2^ = 0.77). The other two alternative ensembles with ordered H12 regions displayed a small but significant decrease in the agreement to the experimental curve, with χ^2^ of 1.37 and 2.44 for the antagonist and agonist positions, respectively (Fig. S1). These results indicate that in the unliganded form, H12 helices of both RXR and RAR are disordered, and highlight the sensitivity of SAXS measurements and analysis to minute structural changes in the heterodimer.

We have exploited this sensitivity to monitor the structural changes in RXR/RAR induced by the binding of two selective RAR ligands, BMS493 (RAR inverse agonist) and Am580 (RAR agonist). The presence of these two ligands induces subtle but noticeable differences in the resulting curves (Fig. 1, Fig. S1 and Table S1). In the presence of BMS493, the observed *R_g_*, 26.5 ± 0.3 Å, is similar to the one measured for the unliganded RXR/RAR heterodimer. Conversely, the presence of Am580 induces a compaction of the particle, with a *R_g_* of 25.6 ± 0.2 Å. We have used the ensemble models based on the available X-ray structures to understand the structural bases of these differences. Concretely, two ensembles of the heterodimer were built in which RXR H12 was maintained disordered whereas RAR H12 was assumed disordered or placed in agonist position (see methods section). The average curves from these ensembles were linearly combined with that of the disordered H12 RAR to optimally describe the measured SAXS curves. SAXS curve of RXR/RAR in the presence of BMS493 is nicely described (χ^2^ = 0.87) with models consisting in a major contribution (85%) of a fully disordered RAR H12 helix (Fig. 1). Conversely, in the presence of Am580, SAXS data are in agreement with the 100% of RAR H12 folded in the agonist position (χ^2^ = 0.88) (Fig. 1).

**Figure 1:**
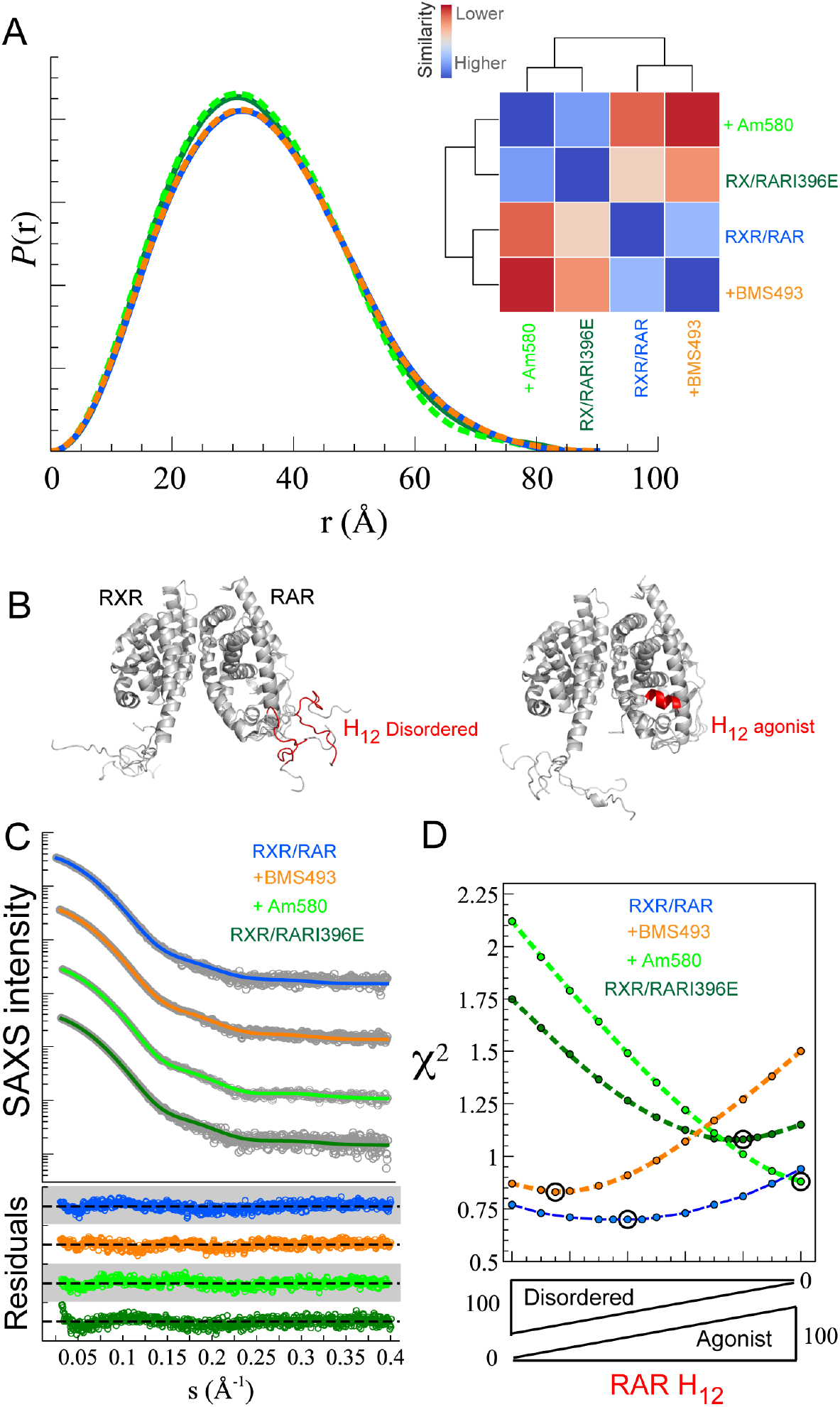
**Modulation of RAR H12 disorder**. (A) Pairwise distance distribution functions, *P(r)*, computed from experimental SAXS curves of wild-type RXR/RAR heterodimer (blue), in the presence of the ligands BMS493 (orange) and Am580 (light green), and for the RXR/RARI396E mutant (dark green). The inset is a heat-map showing their pairwise Kolmogorov-Smirnov-based similarities (KS). The scale is relative from low (red) to high (blue) KS values. (B) Ensemble models (5 conformations) of RXR/RAR with RXR disordered and either RAR H12 in disordered (left) or agonistic (right) conformation. (C) SAXS intensity profiles (gray dots) in logarithmic scale as a function of the momentum transfer s = (4π sin θ/λ, where 2θ is the scattering angle, compared with the optimal combination of conformational ensembles for the RXR/RAR (blue), upon the addition of BMS493 (orange), Am580 (light green), and for the RXR/RARI396E (dark green). Bottom, point-by-point residuals of the fitting with the same color code. (D) Evolution of χ^2^ as a function of the relative contribution of the ensembles corresponding to RXR/RAR with disordered H12 and RAR H12 in an agonistic conformation. Well-defined minima are observed for all scenarios.

Similarly, we have analyzed the conformational changes of RAR H12 helix in the RARI396E point mutant in the context of the heterodimer (RXR/RARI396E). This point mutation, which was designed based on the crystal structure of the complex between RARα and a corepressor peptide, is expected to destabilize the RAR β-strand S3 conformation and favor a helical conformation in H11, thus mimicking agonist-induced conformational change (Le Maire et al., 2010). Interestingly, the SAXS analysis indicates that the main heterodimeric species of this mutant has the RAR H12 in a compact agonistic disposition although a certain population (~20%) of disordered H12 was also observed (χ^2^ = 1.11, Fig. 1D). Not surprisingly, the *P(r)* functions of the mutant RXR/RARI396E and Am580-bound heterodimer are similar, whereas the unliganded RXR/RAR resembles more the BMS493-bound RXR/RAR (Fig. 1A).

In summary, our structural analysis shows that H12 helices of unliganded RXR/RAR heterodimer are mainly disordered in solution, and that the flexibility of RAR H12 can be modulated by the presence of specific ligands or mutation. These observations substantiate the pivotal role of H12 region as a modulator of RAR activity.

### N-CoR_NID_ is a disordered protein with evolutionarily conserved and partially structured elements

We produced and characterized a large fragment of the nuclear receptor corepressor N-CoR (Hörlein et al., 1995) spanning from residue Gln2059 to Glu2325. This fragment, N-CoR_NID_ from now on, corresponds to the nuclear interaction domain (NID) of N-CoR and encompasses the two nuclear receptor binding motifs involved in the interaction of the protein with RXR/RAR heterodimer, CoRNR1 (from 2065 to 2088) and CoRNR2 (from 2269 to 2291) (Fig. 2). The biophysical characterization of N-CoR_NID_ unambiguously indicates that the protein behaves as an Intrinsically Disordered Protein (IDP) (Table S3, Fig. S2 and Fig. 2). Concretely, N-CoR_NID_ displays a reduced ^1^H dispersion in NMR spectra (Fig. 2A), and the Kratky plot does not present a clear maximum (Fig. 2B). Moreover, the *s*_20w_ and frictional ratio measured by SV-AUC, 2.37 ± 0.19 S and 1.53 ± 0.13, are not compatible with a globular protein of this size (≈29 kDa) (Fig. S2). Interestingly, far-UV Circular Dichroism (CD) measured on N-CoR_NID_ presents features that suggest the presence of helical regions (Fig. 2C). Secondary structure is manifested by a shift in the negative maximum at 205 nm, rather than at 198 nm for a pure random coil profile, the negative shoulder near 220 nm, which is more pronounced than that observed for fully disordered proteins, and the positive signal at 190 nm (Bienkiewicz et al., 2002).

We performed the NMR study of N-CoR_NID_ to identify structural features at the residue level (Fig. 2A). The resonances of backbone nuclei of N-CoR_NID_ were assigned using standard triple resonance spectra at high spectrometer field. Out of the 241 expected ^1^H-^15^N HSQC backbone correlation peaks, only 183 could be unambiguously assigned (Fig. S3). The non-identified correlations were either non-visible peaks or had extremely low intensities precluding assignment. When mapping the missing peaks on the amino acid sequence of N-CoR_NID_, they clustered in three non-consecutive regions of the protein (Fig. 2E and Fig. S3B). Interestingly, two of these clusters were centered in the consensus NR binding domains CoRNR1 and CoRNR2 and extended towards both flanking regions (Fig. 2E). Furthermore, a third region, spanning from 2204 to 2234, also displayed absence or systematic decrease of NMR intensities. Within this third region, which will be named Intermediate Region (IR) from now on, no correlation peaks could be assigned for the residues 2204-2217 and 2222-2223. We attributed the absence or decrease of NMR intensities in these three regions to the formation of transient secondary structural elements that experience chemical exchange processes in the *µs-ms* time-scale inducing severe broadening of the signals. The presence of partially structured regions was substantiated by the analysis of N-CoR_NID_ sequence using several disorder prediction servers (Fig. 2F). All predictors applied coincided in identifying CoRNR1 and CoRNR2 as helices and their respective flanking regions as partially structured, both motifs named as ID1 and ID2, respectively, from now on. Interestingly, disorder predictors also identify the IR as partially structured, although to a lesser extent (Buchan et al., 2013).

A sequence conservation bioinformatics analysis of N-CoR_NID_ fragments from multiple eukaryotic organisms indicates that ID1 and the large C-terminal region (LCR, 2190 to 2295), which encompasses IR and ID2, are evolutionarily conserved (Fig. 2G). On the contrary, the N-terminal region of the N-CoR_NID_, with the exception of ID1, is poorly conserved as typically observed in IDPs (Ota & Fukuchi, 2017). Interestingly, the LCR presents a sequence composition that is closer to globular proteins according to the charge hydropathy plot (Uversky et al., 2000), a behaviour that is different from the N-terminus (Fig. 2D).

**Figure 2:**
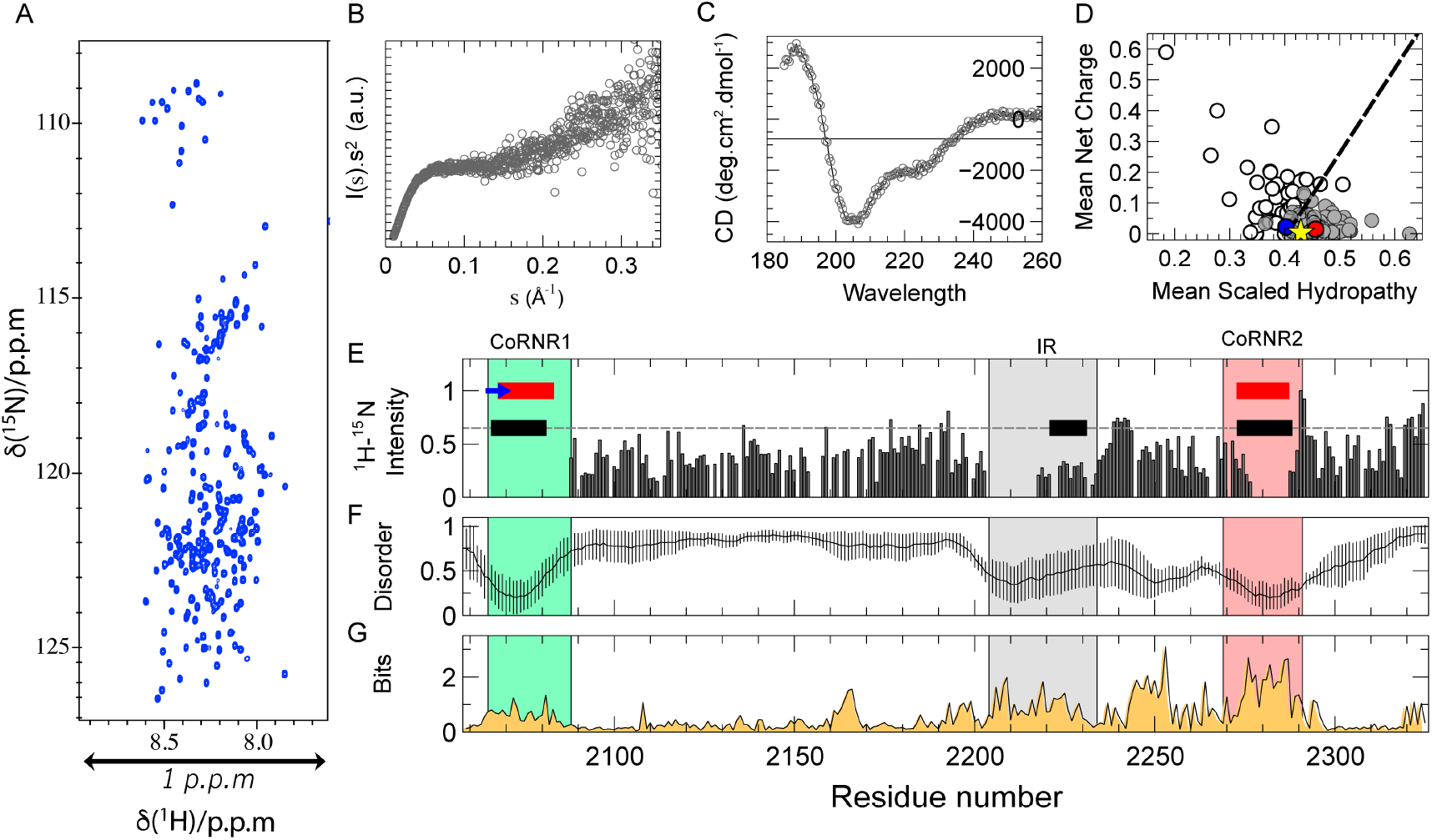
**Structural characterization of N-CoR_NID_**. (A) ^1^H-^15^N HSQC of N-CoR_NID_ displaying the reduced backbone amide ^1^H frequency dispersion typical from IDPs. (B) Kratky representation of the SAXS profile measured for N-CoR_NID_ indicating its lack of compactness. (C) Far UV Circular Dichroism of N-CoR_NID_. (D) Charge Hydropathy (Uverky’s plot) of N-CoR_NID_. Mean net charge versus mean hydropathy plot for the set of 275 folded (grey circles) and 91 natively unfolded proteins (white circles). The N-terminus of N-CoR_NID_ (blue) is clearly in the cluster of disordered proteins, whereas C-terminus of N-CoR_NID_ (red) is on the of folded proteins side. (E) Normalized ^1^H-^15^N HSQC’s intensities of ^15^N -labeled N-CoR_NID_ along the primary sequence (700 MHz, 293 K). The colored bands represent the CoRNR1 and CoRNR2 motifs in green and pink, respectively. The intermediate region (IR) is colored in grey. Secondary structure elements of CoRNR boxes are plotted as a red cylinder and blue arrow for α-helix and β-strand, respectively. The PSIPRED predicted secondary structure elements are represented in black and dashed line for coil. (F) Average disorder prediction (solid grey line) and its standard deviation computed using different computational tools (IUPRED, PrDOS, PONDR-FIT and DISOPRED3) along the N-CoR_NID_ primary sequence. (G) Sequence conservation profile computed from a Hidden Markov Model multiple sequence alignment using Skylign. The overall height indicates the conservation per position.

### N-CoR_NID_ presents intramolecular transient contacts between conserved, co-evolved and partially structured IR and ID2 regions

The presence of local compaction and long-range contacts in the partially structured and conserved C-terminal region of N-CoR_NID_ was explored using SAXS, Paramagnetic Relaxation Enhancement (PRE) NMR experiments, which report on distance-depending induced relaxation on NMR active nuclei, and Molecular Dynamics simulations (MD). In addition to the native Cys2074, which sits in the middle of the ID1 motif, Ser2213 (preceding the IR) and Ser2288 (succeeding ID2) were mutated to cysteines. Note that for these two mutants, the native Cys2074 was mutated into serine (C2074S) to have only one cysteine at a time in the entire sequence of N-CoR_NID_. After incorporating a PROXYL stable radical on each of the cysteine residues independently, PRE-ratios were measured for the three samples. When the paramagnetic moiety was introduced in native Cys2074, no distal effects were observed, indicating that the N-terminal region of N-CoR_NID_ does not present long-range interactions with the rest of the protein (Fig. 3A). Conversely, PRE data measured in the two point mutants, S2213C and S2288C, provide a different picture of N-CoR_NID_. For both single-cysteine variants, a substantial decrease in intensity is observed for ^1^H-^15^N HSQC peaks from the region between the IR and ID2 indicating the presence of extensive long-range contacts (Fig. 3A). Interestingly, PREs measured in this region display a bell-shape with stronger PRE effects in the proximity of both partially structured regions. Moreover, PRE profiles for both mutants in the LCR are very similar suggesting a direct interaction between the IR and the ID2 motifs that also affects the connecting region (Fig. 3A). The compactness of N-CoR_NID_ observed by PRE was substantiated by SAXS. A simple random coil model built with Flexible-Meccano (Bernado et al., 2005; Ozenne et al., 2012) could not reproduce the experimental SAXS curve (Fig. 3B). Indeed, the theoretical ensemble turned out to be more extended than N-CoR_NID_ in solution, with *R_g_* of 50.4 Å and 47.2 ± 1.2 Å for the theoretical and experimental SAXS curves, respectively.

PRE and SAXS data were used to further characterize the structural compaction observed in N-CoR_NID_. Large ensembles of conformations were built with Flexible-Meccano to which multiple explicit dispositions of the PROXYL moiety were attached to the native or engineered cysteine residues. Subsequently, the theoretical averaged PRE ratios were computed as previously described (Salmon et al., 2010), and compared with the three experimental PRE profiles (see methods for details). Not surprisingly, the random coil model only presented contacts in the vicinity of the paramagnetic sites, and therefore did not reproduce the experimental profiles especially in the C-terminal region (Fig. 3A, blue lines). In order to interpret the long-range contacts observed in the LCR, a structurally biased model was built based on previous experimental and bioinformatics observations. Concretely, conformations with at least two contacts of ≤ 15 Å between residues from distal regions of the LCR presenting partial structuration (average disorder below 0.5, Fig. 2F), evolutionary conservation (Bits ≥ 1.6, Fig. 2G), or co-evolution (Fig. 3D), were selected from a large ensemble of random coil conformations. This filtered ensemble resulted too compact, and it was further refined using the SAXS curve to yield a sub-ensemble compatible with the scattering profile (Fig. 3B, C), which was subsequently used to compute the PRE values for the three PROXYL-tagged N-CoR_NID_ constructs and compared with the experimental ones. The resulting theoretical PRE profiles displayed an excellent agreement with the experimental ones (Fig. 3A, red lines). Not surprisingly, a systematic decrease of the PRE values in the LCR compared to the N-terminal region was observed. More interestingly, PRE fluctuations in the connecting region between the IR and the ID2 region were nicely reproduced for the S2213C and S2288C mutants.

**Figure 3:**
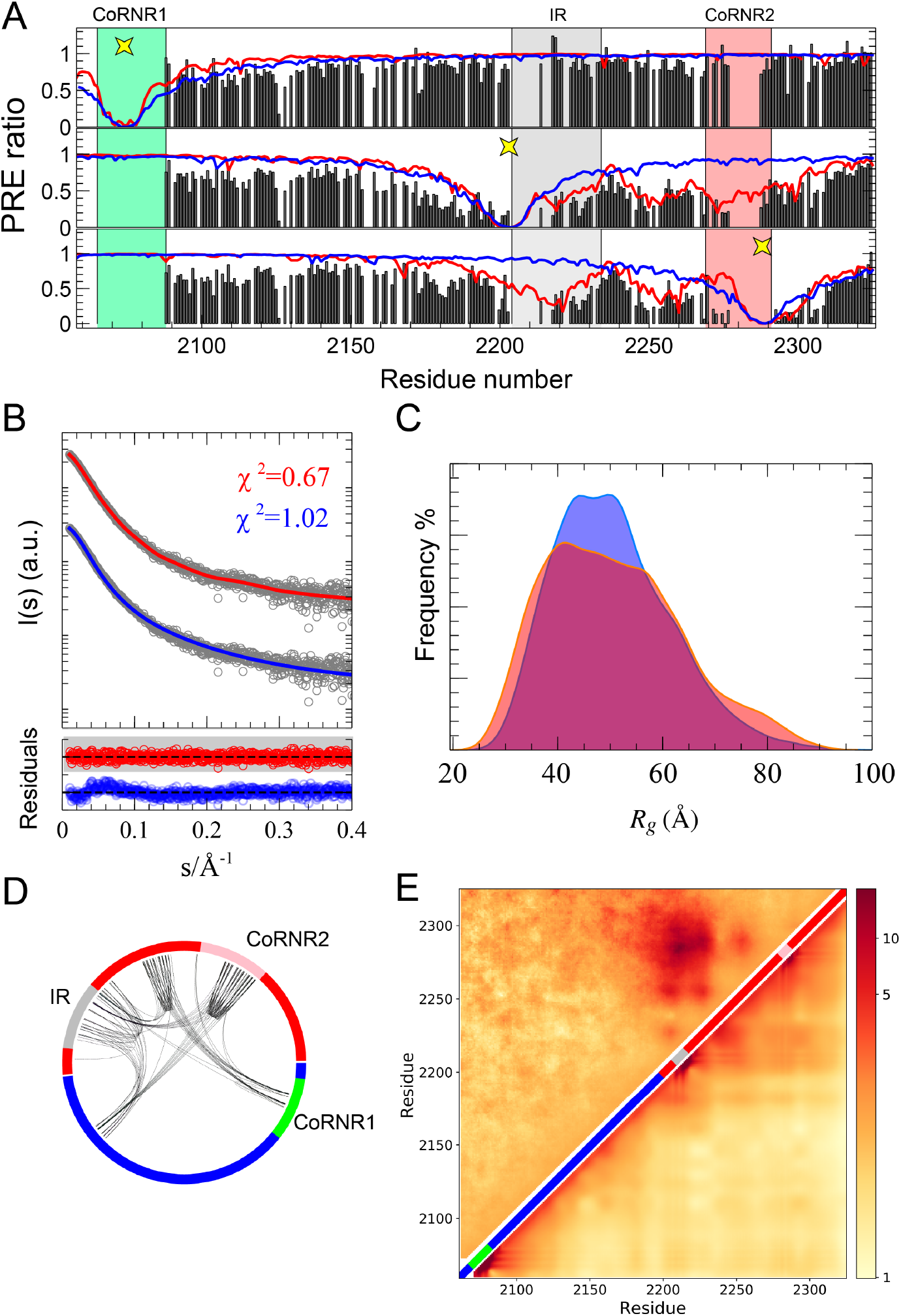
**Long-Range interactions in NCoR_NID_**. (A) Paramagnetic Relaxation Enhancement data of N-CoR_NID_ measured on WT C2074 (top), S2213C (middle) and S2288C (bottom) N-CoR_NID_ mutants. Experimental values (black bars) calculated from the ratio of intensity (PRE Ratio = *I_para_/I_dia_*) of ^1^H-^15^N HSQC spectra measured on the paramagnetic (*I_para_*) samples in the position indicated with a yellow star, and on the diamagnetic samples (*I_dia_*). Averaged theoretical PRE-ratio profiles computed from 10,000 N-CoR_NID_ structures modelled as fully random coil (solid blue line) or assuming long-range interactions (solid red line) between predicted co-evolved, conserved and structured residues within the C-terminal region (LRC). (B) Experimental SAXS curve of N-CoR_NID_ (gray open circles) overlapped with curves calculated form the random coil pool (blue) and EOM selected sub-ensemble with long-range contacts (red). The bottom panel shows the point-by-point residuals of the fitting for both conformational ensembles. (C) Radius of gyration, *R_g_*, distributions obtained for the random coil ensemble (blue) and the EOM selected sub-ensemble (red). (D) Chord-plot of co-evolved residues predicted using BISAnalyser where the lines connect residues identified as co-evolving pairs. (E) Cα-Cα contact map with respect to a random coil model in logarithmic scale computed from the PRE/SAXS refined ensemble (top triangle) and from the coarse-grained MD trajectory (bottom triangle). Color scale quantifies the intensity of the contact where red indicates and average close contact and yellow indicates an average distance equivalent to the random coil model. CoRNR1, IR, and CoRNR2 are displayed in the diagonal to facilitate interpretation of the map.

In order to further investigate the formation of these long-range contacts, we performed MD simulations of N-CoR_NID_ based on a coarse-grained model that was specifically designed for IDPs and takes into account sequence-specific electrostatic and hydrophobic interactions. The resulting conformational ensemble, which was not biased by experimental data, revealed that N-CoR_NID_ sequence is poised to form transient yet noticeable interactions in the C-terminal region. Importantly, the contact matrix derived from coarse-grained MD is in good agreement with the PRE-derived ensemble (Fig. 3E). These results substantiate the presence of transient intramolecular contacts between distal regions within the C-terminal region of N-CoR_NID_. Moreover, these regions present relatively stable secondary structural elements and a high level of evolutionary conservation.

### RXR/RAR heterodimer interacts with the corepressor mainly through RAR and in a cooperative manner

After characterizing the individual partners, we investigated the complex between RXR/RAR and N-CoR_NID_. The affinities of RXR/RAR heterodimer for NCoR peptides encompassing either CoRNR1 or CoRNR2 motifs and for the N-CoR_NID_ fragment were measured by fluorescence anisotropy and thermophoresis, respectively. As shown in Figure 4, formation of the RXR/RAR heterodimer does not modify the binding capacity of individual RXR and RAR monomers for isolated CoRNR peptides. In fact, the RXR/RAR heterodimer binds the two isolated CoRNR peptides with nearly the same affinity, 1.68 and 1.47 µM for CoRNR1 and CoRNR2, respectively (Fig. 4B). These affinity values are very similar to those measured previously (Le Maire et al., 2010) for the unliganded RAR monomer (1.40 and 1.55 µM for peptides encompassing CoRNR1 and CoRNR2, respectively) (Fig. 4A). The slight preference observed for CoRNR1 is induced by the ability of this motif to form a β-sheet interface with S3 of RARα, which is further stabilized in the presence of the inverse agonist BMS493 with an affinity of 0.17 µM. Conversely, in its monomeric form, RXR presents a moderate affinity (22.5 µM) for CoRNR2, and no measurable interaction with CoRNR1 (Fig. 4A). As expected, the presence of the RAR-selective agonist Am580 or the RARI396E mutation cause a noticeable decrease in the affinity of RXR/RAR for both N-CoR peptides with a stronger effect on CoRNR1 (Fig. 4B). Conversely, the RAR inverse agonist BMS493 efficiently increases the binding affinity of RXR/RAR for CoRNR1, but has not much effect on the interaction with CoRNR2 (Fig. 4A-B). These observations suggest a sequential and directional mechanism in which, in the context of N-CoR_NID_, comprising the two CoRNR motifs, the main anchoring point would involve RAR on the heterodimer side and CoRNR1 on the N-CoR side. This primary contact would then enable a second lower-affinity interaction between RXR and CoRNR2.

Microscale thermophoresis measurements further substantiated the cooperativity and directionality of the interaction. The affinity of RXR/RAR heterodimer for N-CoR_NID_ was found to be much higher than for individual peptides with a value of 0.21 ± 0.09 µM, to be compared to 1.68 and 1.47 µM for CoRNR1 and CoRNR2, respectively (Fig. 4B-C). The inverse agonist BMS493 or deletion of RXR helix H12 (RXRΔH12), which are known to enhance the interaction of RAR with CoRNR1 (Germain et al., 2009; Le Maire et al., 2010) (Fig. 3A) and that of RXR for CoRNR2 (Hu and Lazar, 1999), respectively, were shown to increase significantly the overall affinity of the heterodimer for N-CoR_NID_ (Fig. 4C). In contrast, the RAR agonist Am580, or the RAR mutation I396E induced a strong decrease of the affinities with Kd values of 8.6 ± 3.70 µM or 4.42 ± 1.86 µM, respectively, due to the weakening of the interaction of RAR with CoRNR1 in both cases. The observation that the perturbation of individual anchoring points has severe consequences on the affinity of RXR/RAR for N-CoR_NID_ demonstrates the cooperativity and directionality of the complex.

**Figure 4:**
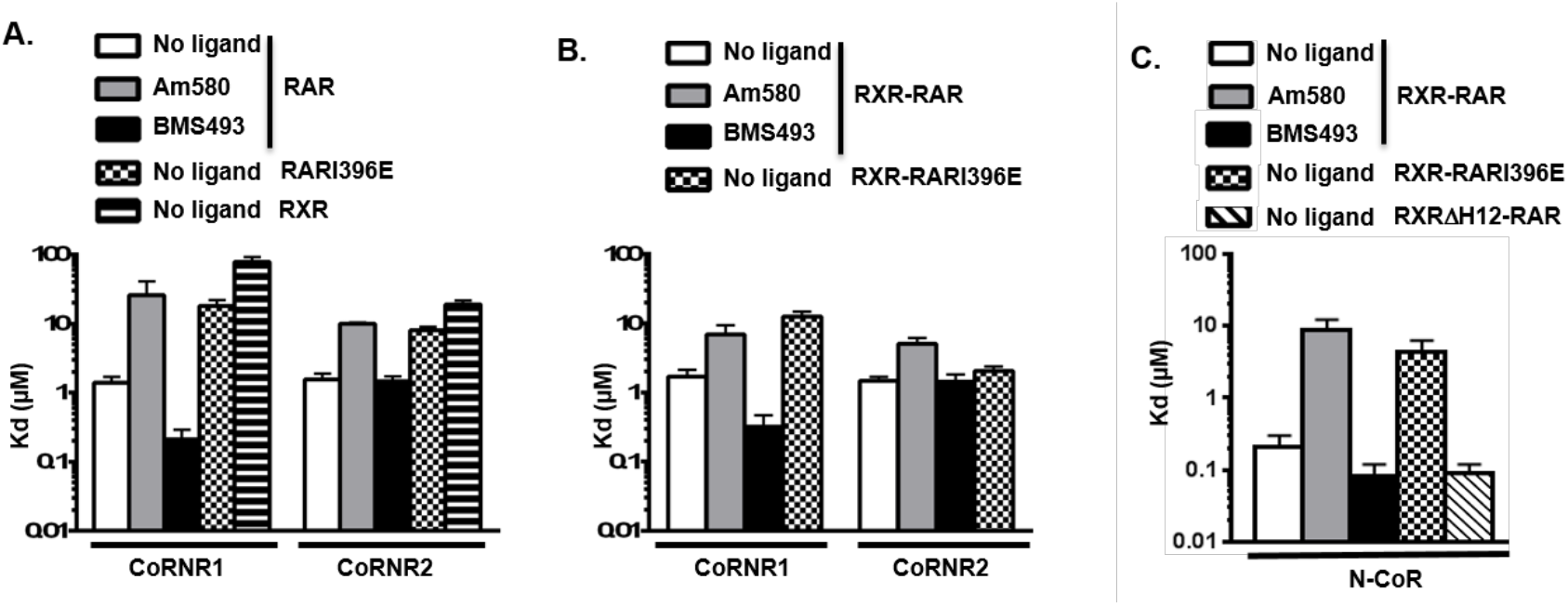
**RXR/RAR heterodimer interacts with N-CoR_NID_ mainly through RAR**. (A) Affinities of CoRNR motifs of N-CoR for wild-type RAR, mutant RARI396E and RXR measured by fluorescence anisotropy, in the absence of ligand or the presence of Am580 (RAR agonist) or BMS493 (RAR inverse agonist). Data on RAR and RARI396E were previously published in (Le Maire et al., 2010) and are added here for clarity. (B) Affinities of CoRNR motifs of N-CoR for wild-type RXR/RAR and mutant RXR/RARI396E heterodimers measured by fluorescence anisotropy in same ligation states as in A. (C) Affinities of N-CoR_NID_ fragment (comprising both CoRNR1 and CoRNR2 motifs) for wild-type RXR/RAR (in same ligation states as in A), mutant RXR/RARI396E and RXRΔH12/RAR heterodimers measured by Microscale Thermophoresis.

### N-CoR_NID_ forms a transient multi-site complex with RXR/RAR

Affinity measurements indicate that N-CoR_NID_ binds cooperatively to RXR/RAR heterodimer. To unveil the structural bases of this cooperativity, we combined SAXS and NMR to study the N-CoR_NID_:RXR/RAR heterotrimeric complex.

SAXS data indicates that the complex is monodisperse with a *R_g_* and *D_max_* values of 48.4 ± 1.1 Å and 194 ± 10 Å, and with a stoichiometry of 1:1:1 according to SV-AUC and mass spectrometry data (Table S3, Fig. S2 and Fig. S4). Moreover, the Kratky representation with elevated baseline at high *s*, and asymmetric *P*(r) function suggest that N-CoR_NID_:RXR/RAR has a significant degree of flexibility (Fig. S5).

The observation of cooperative effects prompted towards the possibility that the two binding regions of N-CoR_NID_, CoRNR1 and CoRNR2, could simultaneously interact with RXR/RAR forming the so-called deck model (le Maire and Bourguet, 2014). The latter scenario is in contrast with the asymmetric model, where a single corepressor binding side would interact with the RXR/RAR heterodimer as observed for the complex between RXR/RAR and a fragment of the coactivator Med1 (Rochel et al., 2011). Ensembles of conformations were built for both the deck and the asymmetric binding modes (see methods section for details), integrating previous knowledge of the system. This includes the directionality of the complex derived from affinity measurements, and two crystallographic structures of RAR and RXR with corepressor peptides that were used to define the atomic details of the interacting sites (Table S2) (Le Maire et al., 2010; Zhang et al., 2011). H12 helices from both RXR and RAR were considered disordered based on our SAXS analysis (see above). Finally, the non-interacting regions of N-CoR_NID_ were assumed to adopt random coil conformations based on the NMR data (see below). Theoretical SAXS curves were computed from large ensembles of structures for both interacting modes yielding notably different profiles (Fig. S6). These curves were optimally combined to derive the relative population of both interacting modes by minimizing the agreement to the experimental curve. An excellent agreement (χ_i_^2^ = 0.88) to the experimental SAXS curve of N-CoR_NID_:RXR/RAR was obtained when a 85/15 ratio for the asymmetric/deck models was used (Fig. 5A, B). This result indicates that both binding modes coexist in solution although the asymmetric model is the major species in the absence of ligand. The similarity of the scattering curves between the two alternative asymmetric models (N-CoR_NID_ bound to RAR through ID1 or N-CoR_NID_ bound to RXR through ID2) precludes the identification of the major asymmetric species. However, the higher affinity of ID1 for RAR suggests that the former is the dominant docking side in the asymmetric model.

In order to understand the interaction at the residue level, we performed NMR experiments by adding an equimolar amount of RXR/RAR to a ^15^N-isotopically labeled N-CoR_NID_ sample. The NMR ^1^H-^15^N HSQC spectrum of N-CoR_NID_ presents correlation peaks that are superimposable to these obtained in the absence of heterodimer, and no correlation peaks shifted upon the addition of the heterodimer (Fig. S7). This observation indicates that the conformational properties of N-CoR_NID_ are equivalent in the free and in the bound states, and that N-CoR_NID_ remains mostly disordered when bound to the RXR/RAR heterodimer. However, in the presence of RXR/RAR, a systematic decrease in peak intensities is observed for N-CoR_NID_ when compared to the free form (Fig. S7 and Fig. 2E). We attribute this observation to the perturbation of the hydrodynamic properties of N-CoR_NID_ upon binding to the heterodimer that senses the presence of a large globular particle and increases the apparent correlation time. Interestingly, the intensity decrease is not homogeneously distributed along the protein, and two regions of the protein can be clearly distinguished. The N-terminal region connecting ID1 and the IR presents moderate intensity decreases that are slightly larger in the proximity of both partially structured regions. Conversely, the region connecting the IR with ID2 displays a more important intensity reduction. The transient binding of ID2 to RXR as demonstrated by SAXS and the change in the dynamic regime for the intra-molecular interactions could explain the enhanced intensity reduction in the C-terminus. To explore this phenomenon, we performed PRE experiments for the complexes formed by the previously described N-CoR_NID_ cysteine mutants and the heterodimer RXR/RAR. The incorporation of radical moieties on the N-CoR_NID_ S2213C and S2288C point mutants induced a sequence-dependent decrease on the PRE ratios of residues placed at their flanking regions, a phenomenon which is typically observed in IDPs (Fig. 5D, E). Interestingly, for both mutants a small but systematic increase in the PRE ratios was found for the N-terminal residues when compared with the PRE data of the free forms (Fig. 3A). This is most probably caused by the reduced conformational exploration in N-CoR_NID_ when bound to the heterodimer, which limits sporadic contacts of the C-terminal region with the N-terminus. Compared with the free form of N-CoR_NID_, the presence of the heterodimer causes a PRE-ratio increase in the region connecting the IR and ID2, and partially suppresses the above-described bell-shape of PRE ratios. This increase is not homogeneous and the region around 2250 presents strong PRE effects. These observations indicate that the population of conformations experiencing long-range contacts in the C-terminus is diminished in the bound form. This last observation is coherent with the existence of a minor population of the deck model that conformationally restricts N-CoR_NID_, and partially hampers intra-molecular contacts involving ID2.

**Figure 5:**
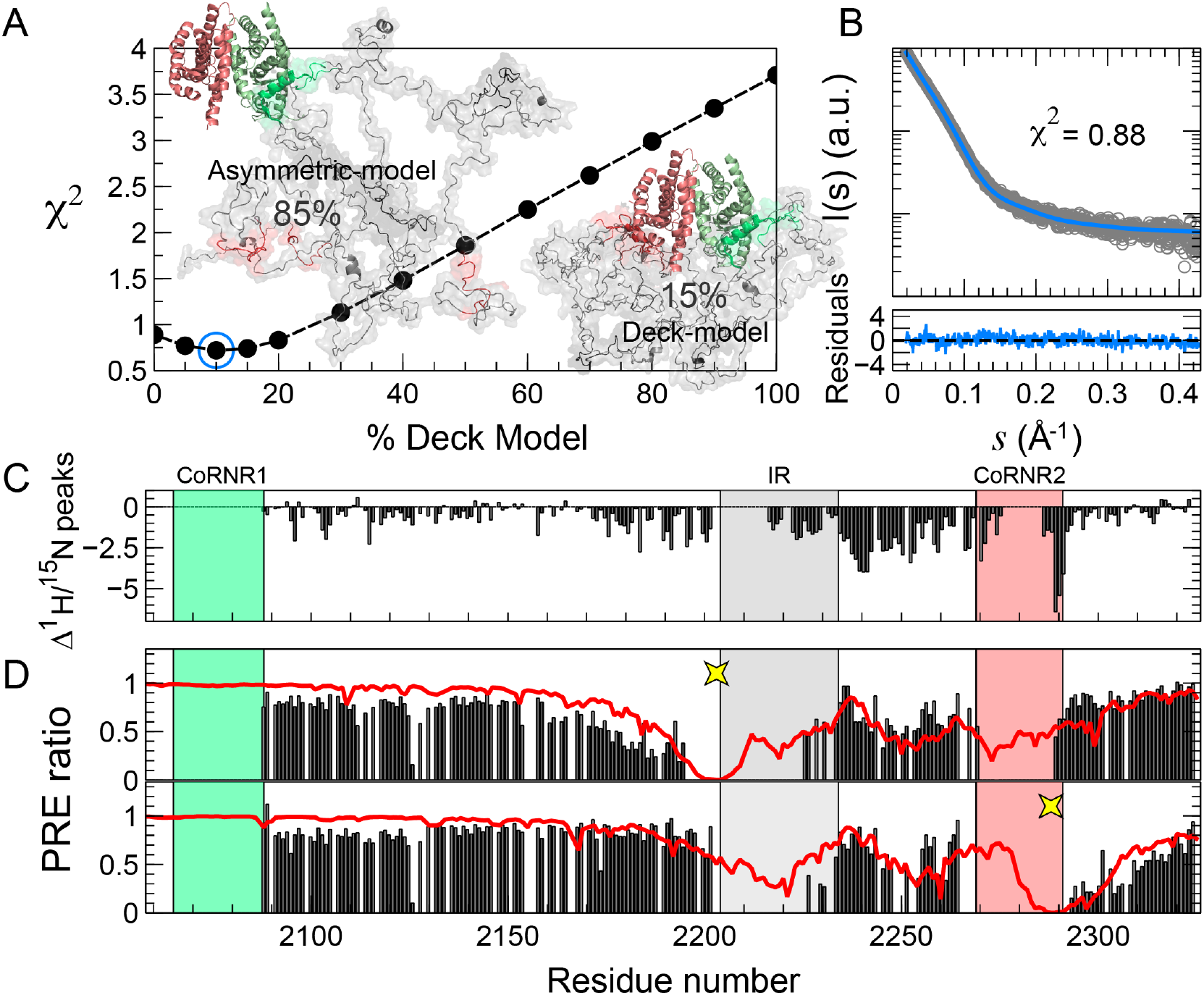
**Structural bases of the cooperative binding of N-CoR_NID_ with the RXR/RAR heterodimer**. (A) Optimization of the relative population of the asymmetric and deck forms of the N-CoR_NID_:RXR/RAR. χ^2^ values (black dots) for different relative populations of the two forms of the complex. Ten conformations for each of the ensembles, asymmetric and deck models, are shown. (B) Scattering intensity, *I(s)*, as a function of the momentum transfer, *s*, for the SAXS curve measured for N-CoR_NID_:RXR/RAR (grey empty dots) and the theoretical one for the combination of the asymmetric and deck models (85:15) (blue line). Residuals are displayed at the bottom of the figure to substantiate the quality of the agreement. (C) ^1^H-^15^N HSQC peak intensity differences for N-CoR_NID_ in its free state and in the presence of an equimolar amount of RXR/RAR. (D) PRE values for S2213C (top) and S2288C (bottom) N-CoR_NID_ mutants (positions of the paramagnetic tags are indicated with yellow stars) in the presence of the RXR/RAR heterodimer (black bars). Theoretical PRE values obtained for a structurally biased model of free N-CoR_NID_ (equivalent to Fig. 3A) are displayed to highlight the differences in the long-range intramolecular interaction network between the free and bound states of the protein.

### Cognate ligands and mutations modulate the conformational equilibrium in the complex

Thermophoresis experiments demonstrated that the affinity of N-CoR_NID_ for RXR/RAR can be finely tuned by the addition of cognate ligands or by mutations at the recognition sites of both RXR and RAR. We applied SAXS and NMR to explore the structural bases of this affinity modulation.

The addition of the RAR inverse agonist BMS493 reinforced the interaction of RXR/RAR with N-CoR_NID_ through CoRNR1 (Fig. 4B and 4C). The analysis of the SAXS curve of the ternary complex in the presence of BMS493 indicates that the overall size of the particle is slightly reduced with respect to the unliganded form of the complex, *R_g_* = 47.5 ± 1.0Å (Table S3). The characterization of that curve in terms of atomistic ensemble models indicated that the asymmetric model is still the major species but the population of the doubly-bound deck model increases up to 35% (Fig. 6A and 6B). A more extreme situation was observed when the complex was formed with RXRΔH12/RAR that strongly reinforces the overall affinity with N-CoR_NID_ by increasing the interaction of RXR with ID2 (Fig. 4B and 4C). This RXR H12 deletion renders the RXR hydrophobic groove more accessible and has been reported to significantly increase the interaction of RXR with corepressor (Hu and Lazar, 1999; Schulman et al., 1996; Zhang et al., 2011). In that situation, the SAXS curve, which presented a smaller *R_g_* of 42.0 ± 2 Å, could be explained with the only presence of the deck model (Table S3, Fig. S5 and Fig. 6A and 6B). The ^15^N N-CoR_NID_ NMR intensities measured for both complexes, upon addition of RXR/RAR in the presence of BMS493 and of RXRΔH12/RAR, displayed a systematic decrease with respect to the unliganded N-CoR_NID_:RXR/RAR (Fig. 6C). We attributed this observation to the increase of the overall correlation time sensed by N-CoR_NID_ when the population of the deck model increases. This is especially significant in the region close to ID2 that fits well with the higher interaction of ID2 with RXRΔH12. Compared with the unliganded (Fig. S7), the region connecting the IR and ID2 in the N-CoR_NID_:RXRΔH12/RAR displays a less prominent decrease of intensities, indicating that some of the structural and/or dynamical properties that cause the decrease in the native complex are abolished by the formation of a doubly-bound state.

We also performed SAXS experiments of the complex N-CoR_NID_:RXR/RAR in the presence of the RAR agonist Am580, and of the complex N-CoR_NID_:RXR/RARI396E. Affinity measurements indicated that both conditions diminished the interaction of RXR/RAR with N-CoR_NID_ through CoRNR1 (Fig. 4B and 4C). The analysis of the SAXS data using explicit models indicated that the decrease of the interaction has severe structural consequences for both complexes. On one hand, the SAXS curve measured on N-CoR_NID_:RXR/RAR in the presence of Am580 can only be described (χ^2^ = 0.81) if large populations, 51%, of unbound N-CoR_NID_ and RXR/RAR are invoked (Fig. 6A and 6B), showing that Am580 partially breaks the complex by diminishing the interaction of CoRNR1 with RAR. Similarly, the SAXS curve measured for the complex N-CoR_NID_:RXR/RARI396E is optimally described using a combination of unbound species, asymmetric complex and a low percentage of deck complex. ^15^N N-CoR_NID_ NMR intensities measured in these two conditions were also coherent with SAXS observations (Fig. 6C). The general increase in these intensities is in line with the presence of a weaker complex with lower dragging forces and smaller apparent correlation time.

**Figure 6:**
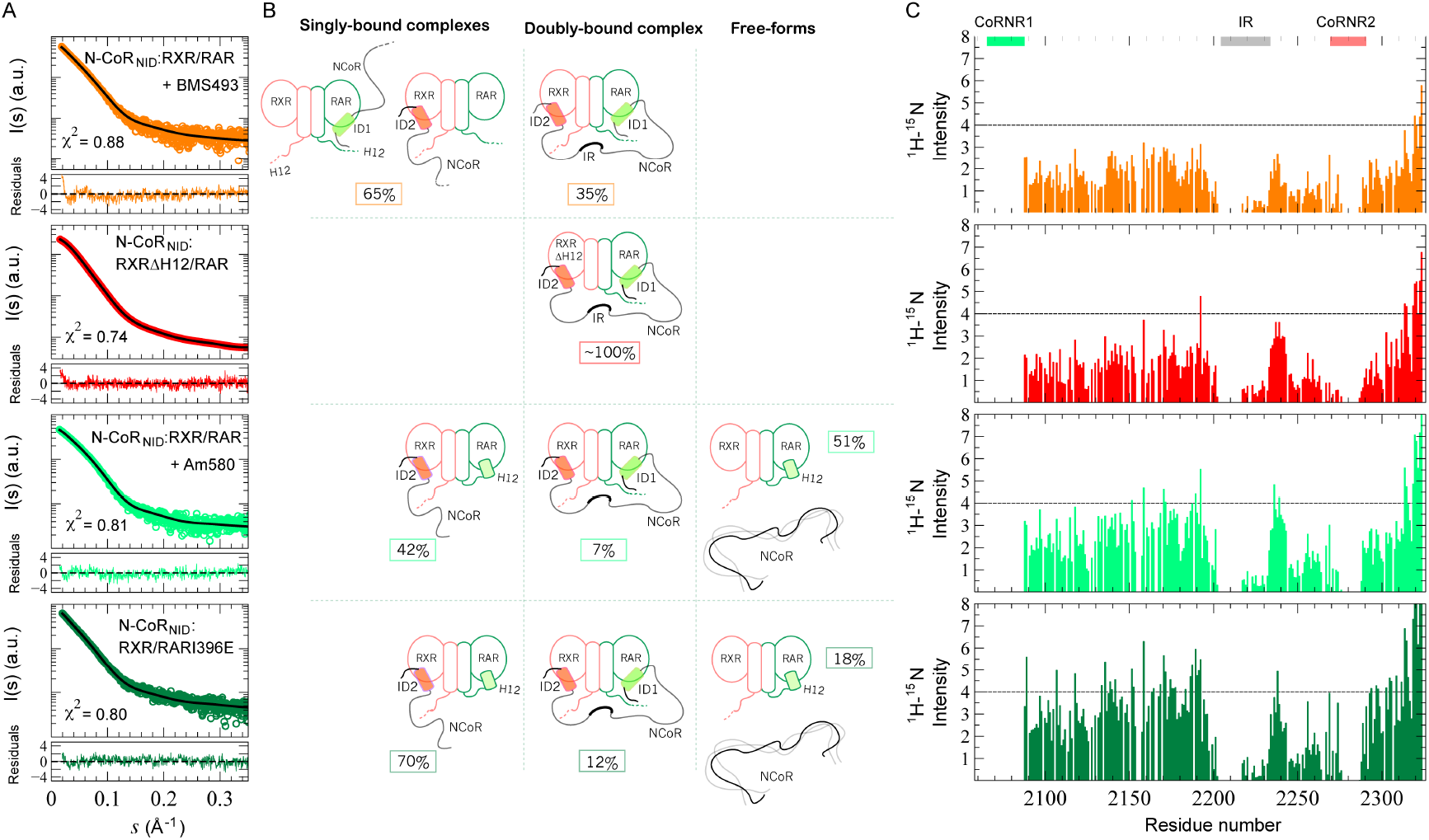
**Modeling of N-CoR_NID_:RXR/RAR complexes**. (A) Logarithmic representation of SAXS intensity, *I(s)*, versus momentum transfer *s* (open circles) of N-CoR_NID_:RXR/RAR in the presence of inverse agonist BMS493 (orange) or agonist Am580 (light green), and N-CoR_NID_: RXRΔH12/RAR (red) and N-CoR_NID_:RXR/RARI39E (green) variants. Simulated scattering patterns (solid black lines) are the best linear combination of theoretical scattering profiles of the species in panel B weighted by their relative population. χ^2^ values indicate the goodness-of-fit. The bottom panels show the point-by-point residuals of the fitting for each scenario. (B) Schematic cartoon representation for all-atom ensembles generated for the different coexisting species in solution. RXR, RAR and N-CoR_NID_ are colored in orange, green and grey, respectively. ID1 (green) and ID2 (red) NR binding motifs are represented as boxes in the NCoR_NID_ cartoon. The relative population of the different species are inside boxes with the same color code as the associated experimental SAXS. (C) ^1^H-^15^N HSQC intensities along N-CoR_NID_ in complex with wt RXR-RAR upon addition of RAR inverse agonist BMS493 (orange), or RAR agonist Am580 (light green); or in complex with RXRΔH12-RAR (red) or RXR/RARI396E (green). The colored bars represent the CoRNR1 and CoRNR2 motifs in green and pink, respectively.

### Mammalian two-hybrid experiments confirm the interaction *in vivo*

Mammalian two-hybrid experiments were performed to validate in a cellular context the cooperativity and the directionality of the interaction between N-CoR_NID_ and RXR/RAR heterodimer, deduced from the measurements of *in vitro* binding constants and from the SAXS/NMR modelling (see above). Two-hybrid analyses were performed in COS cells with chimeras containing GAL4 DNA-binding domain fused to a N-CoR fragment encompassing ID1 and ID2 (from 1629 to 2453 termed Gal-NCoR) or either CoRNR1 or CoRNR2 of N-CoR (Gal-CoRNR1 and Gal-CoRNR2, respectively), and the LBD of RXRα or of RARα fused to the activation domain of VP16 (termed VP16-RXR or VP16-RAR, respectively). Here RAR-selective agonist TTNPB activating RARα as efficiently as Am580 was used as the reference.

First, we confirmed that wild-type RXR/RAR heterodimer interact with both CoRNR1 and CoRNR2 (Fig. 7A), with a slightly better effectiveness for CoRNR1. However, the interaction of RXR/RAR heterodimer with the longer N-CoR fragment (equivalent to N-CoR_NID_) is significantly stronger than the individual CoRNR regions. Therefore, in a cellular environment the two CoRNR motifs in N-CoR bind to the RXR/RAR heterodimer in a cooperative manner and may induce a stronger affinity. In addition, the selective RAR agonist TTNPB is able to release CoRNR1 and the N-CoR fragment from RXR/RAR heterodimer (Fig. 7A). Importantly, a significant interaction of CoRNR2 is measured even in the presence of TTNPB which only targets RAR likely reflecting the interaction of RXR with the CoRNR2 motif (Fig. 7A). The directionality of the interaction between N-CoR_NID_ and RXR/RAR heterodimer was further confirmed with two-hybrid experiment with the N-CoR fragment (Fig. 7B). Whereas RXR alone interacts very weakly with N-CoR, in agreement with previous observations of our group (Germain et al., 2002) and with our fluorescence anisotropy data on RXR monomer and CoRNR peptides (Fig. 4), addition of RARα allows to measure a significant interaction between the RXR/RAR heterodimer and Gal-NCoR. Interestingly, the addition of the RAR-selective agonist TTNPB was sufficient to decrease this interaction (Fig. 7B). Moreover, RAR alone can efficiently recruit N-CoR (Fig. 7C). All these observations confirm that RAR is indispensable for the corepressor recruitment by RXR/RAR heterodimer. On the other side, the role of RXR for the recruitment of N-CoR was subsequently addressed in a cellular environment (Fig. 7C). Relative to RAR alone, addition of RXR yielded a slight but significant increase of the N-CoR interaction confirming an active role of RXR in the heterodimeric form (Fig. 7C), and thus the cooperativity in the interaction. This effect was more pronounced for the RXRΔH12 deletion mutant (Fig. 7C), in agreement with MST experiments and SAXS modelling. We reasoned that the RXR H12 deletion may generate a new interaction surface for N-CoR. Contrary to the wild-type heterodimer, TTNPB does not reduce N-CoR interaction with RXRΔH12/RAR heterodimer (Fig. 7C). This observation indicates that RXRΔH12/RAR retains the ability to interact with N-CoR, through the RXR hydrophobic groove. Two-hybrid experiments with RXRΔH12/RAR heterodimer and Gal-CoRNR1 and Gal-CoRNR2 clearly demonstrated that CoRNR2 interacts with the more accessible groove of RXR as TTNPB was unable to reduce CoRNR2 association, and that CoRNR1 interacts with RAR as its binding is impaired by addition of TTNPB (Fig. 7D), confirming the directionality in the interaction of N-CoR with RXRΔH12/RAR.

**Figure 7:**
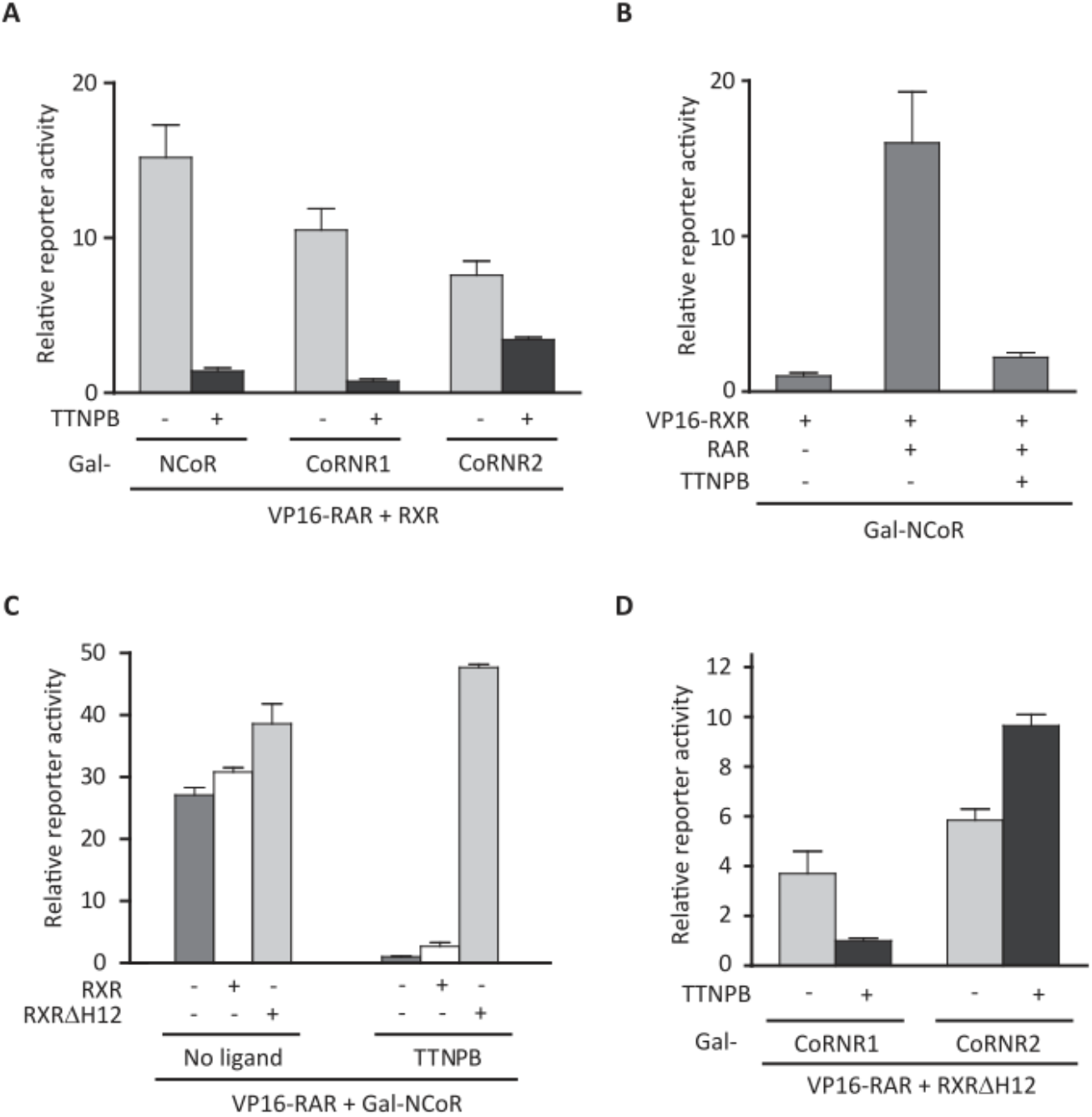
**Interactions of RXR/RAR heterodimers with N-CoR using mammalian two-hybrid assays with (17m)_5x_-G-Luc reporter in COS cells**. (A) Transient transfections with VP16-RAR and RXR, and with Gal-NCoR or Gal-CoRNR1 or Gal-CoRNR2. Both CoRNR1 and CoRNR2 are able to interact with the RXR/RAR heterodimer. (B) Transient transfections with Gal-NCoR and with VP16-RXR alone or in combination with RAR. RXR only minimally interacts with N-CoR and RAR is required for N-CoR-heterodimer complex formation. (C) Transient transfection with Gal-NCoR and with VP16-RAR alone (dark grey) or in combination with RXR (white) or RXRΔH12 (light gray). (D) Transient transfections with VP16-RAR and RXRΔH12, and with Gal-CoRNR1 or Gal-CoRNR2. N-CoR interacts with the RXRΔH12/RAR heterodimer through an interaction between CoRNR2 and the RXR groove. In all assays TTNPB was used at 10nM.

## Discussion

In this study we have integrated multiple structural, biophysical and cell-biology techniques to decipher the molecular bases of transcriptional repression of the retinoic acid nuclear receptor by the corepressor N-CoR. Our results demonstrate that the molecular mechanism relies on the interplay of the flexible elements found in RXR/RAR heterodimer and the intrinsically disordered N-CoR.

We have exploited the high sensitivity of SAXS to probe minute structural changes that perturbe the shape of the RXR/RAR LBDs heterodimer in the absence or in the presence of various RAR ligands and point mutations. The main structural changes of the heterodimer reside in the conformation and position of helices H12 of each monomer. Our analyses demonstrate that helices H12 of RAR and RXR are better described as disordered in the absence of ligand, in line with previous NMR and amide hydrogen/deuterium (H/D) exchange experiments for a number of NRs including RXR (Johnson et al., 2000; Kallenberger et al., 2003; Lu et al., 2006). These results invalidate the initial hypothesis of allosteric inhibition of RXR by a subset of nuclear receptors, such as RAR and THR. It was suggested that, in unliganded RXR/RAR and RXR/THR heterodimers, H12 of RXR docks to the coregulator-interaction site of the partner (Westin et al., 1998; Zhang et al., 1999).

The lack of structural order of RAR helix H12 was also deduced from SAXS data measured on RXR/RAR heterodimer in the presence of RAR inverse agonist (BMS493), in agreement with the absence of density for RAR H12 in the crystal structure of the complex formed by RARα-LBD, the corepressor N-CoRNR1 and the inverse agonist (Le Maire et al., 2010). These two forms of RXR/RAR heterodimers, unliganded and in complex with the inverse agonist, bind corepressors, confirming that the disordered helices H12 are not directly involved in corepressor binding. Nevertheless, the disordered and highly flexible RXR and RAR helices H12 may sterically screen the interaction with corepressors. Indeed, RXRΔH12 and RARΔH12 deletion mutants recruit corepressors more efficiently than wild-type RXR and RAR (Le Maire et al., 2010; Zhang et al., 1999), probably due to a higher accessibility of the LBD interaction region for the corepressors. Conversely, addition of RAR agonist (Am580) induces a compaction of the heterodimer that fits with RAR helix H12 adopting the agonist position, as observed in crystal and solution structures of RAR in the presence of agonists (Bourguet et al., 2000; Egea et al., 2001; Klaholz et al., 2000; Le Maire et al., 2010). Our SAXS analysis of the mutant RARI396E, for which no structural data had been reported before, and whose mutation was suggested to induce the S3 strand to H11 helix transition (Le Maire et al., 2010), also provokes a main rearrangement of RAR helix H12 that is compatible with an agonist conformation. The transition from a disordered state to an ordered and active conformation of RAR helix H12 is induced by the binding of an agonist that efficiently triggers corepressor release. Importantly, this release is dependent of the proportion of agonist conformation of this helix. With this analysis we confirm that the disorder to order transition of helix H12 conformation is a key mechanism of corepressor release and coactivator recruitment by RXR/RAR, contrary to the region corresponding to H11/S3 in RAR which is the master regulator of corepressor association to RAR (Le Maire et al., 2010).

Our extensive experimental and computational analysis of N-CoR_NID_ demonstrates that it is a disordered protein presenting local and long-range structural phenomena that are directly linked to its function. With the exception of the CoRNR1 motif, the N-terminal region presents the prototypical spectroscopic features of a random coil. Moreover, bioinformatics analyses on this region indicate a poor evolutionary conservation and the absence of co-evolutionary interactions with other parts of the protein as often been observed in disordered regions without direct functional roles (Ota and Fukuchi, 2017). Conversely, the C-terminal region including the CoRNR2 motif presents multiple structural features and is evolutionary conserved. In solution, CoRNR1 and CoRNR2 are preformed molecular recognition elements (MOREs) (Mohan et al., 2006) that mediate the interaction with the heterodimer. The lack of NMR information for these regions is probably due to intermediate exchange regime processes linked to the transient formation of secondary structures. Very interestingly, NMR has unveiled a new region, IR, placed between the two NR interaction domains that is partially ordered and highly conserved in eukaryotes. These two observations suggest a relevant functional role for IR. Indeed, N-CoR_NID_ experiences transient long-range tertiary interactions between the IR and ID2 that we have identified and structurally characterized using PREs. The accurate description of experimental PREs was achieved when the long-range interaction between co-evolved residues in partially structured regions of the C-terminal region of N-CoR_NID_ was used. This long range contact is partially impaired when N-CoR interacts with NRs as demonstrated by our NMR analysis of N-CoR_NID_ in complex with the RXR/RAR heterodimer. The formation of the complex would liberate the IR that would become available to interact with other proteins of the repressive macromolecular complex.

A number of structural and biophysical studies have already revealed the complex interactions between NR heterodimers and coactivators (Chandra et al., 2017; Osz et al., 2012; Pavlin et al., 2014; Pogenberg et al., 2005; Rochel et al., 2011; de Vera et al., 2017; Zheng et al., 2017). However, the interaction of NRs with corepressors that hampers gene transcription in the basal state are poorly understood. Our study, which included solution-state structural methods along with biochemical, biophysical and computational approches, reveals for the first time the atomistic details in terms of ensembles of the interaction between RXR/RAR heterodimer and N-CoR. Our study shows a cooperative interaction of both CoRNR1 and CoRNR2 motifs with RXR/RAR as the binding affinity of N-CoR_NID_ is stronger than the individual interactions. In addition, the interaction has a defined directionality: N-CoR_NID_ is recruited primarily to RAR through CoRNR1 enabling CoRNR2 to subsequently bind to RXR for which it has a moderate affinity. Importantly, the other configuration would not produce a cooperative binding as CoRNR1 does not interact with RXR.

**Figure 8:**
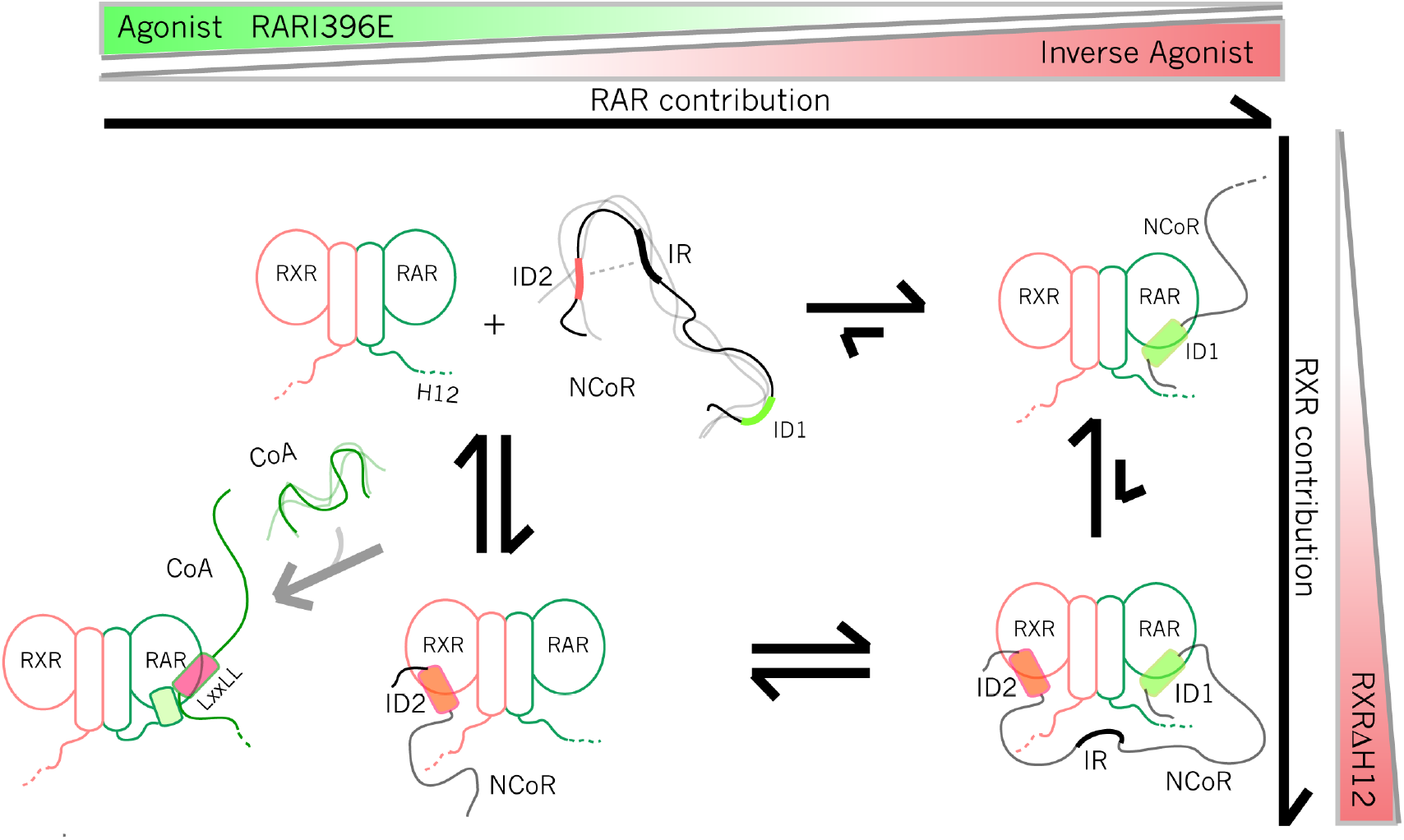
**Schematic illustrations summarizing the structural and thermodynamic data presented in this study**. The central part represents the free partners, RXR/RAR and N-CoR_NID_, and the three forms of the assembly. Size of arrows connecting these states indicates the thermodynamic preferences observed in our study. Equilibrium composition can be modified upon addition of ligands or the presence of mutations on the RXR/RAR heterodimer. These effects are depicted as gradient colored triangles on the top (affecting the RAR binding) and the right (affecting the RXR binding) sides of the figure. Asymmetric form bound through the CoRNR2 and the dissociated RXR/RAR can evolve towards the formation of complex with a coactivator upon addition of an RAR agonist. This transition from a repressive to an activated state of the NR is depicted on the rigth part of the figure.

Taken toghether, our data suggest the mechanistic model that is depicted in Figure 8. In the absence of ligands, it exists an equilibrium between a major population of assymetric binding of N-CoR_NID_ to RXR/RAR and a minor population of a doubly bound N-CoR_NID_ (deck binding mode) in which both CoRNR boxes simultaneously interact with the heterodimer, accounting for the cooperativity of the interaction. The major species of the assymetric binding mode can be resonnably assigned to NCOR_NID_ binding to RAR through the CoRNR1 motif, which presents the strongest local affinity. Our observations *in vivo* through mammalian two hybrid experiments substantiate with this model, also suggesting that the structural phenomena probed *in vitro* also occurs in cells. The addition of RAR ligands or point mutations in either RXR and RAR modify the strenght of the individual interactions and, consequently change the cooperativity and the equilibrium between the assymetric and the deck binding modes. Concretely, the addition of a RAR inverse agonist, which strenghtens the interaction of CoRNR1 with RAR, or the use of a heterodimer with the truncated RXR H12 (RXRΔH12), which increases the interaction of CoRNR2 with RXR, produce a notable increase of the overall affinity of the complex. This increase is a consequence of an enhancement of the cooperativity by the equilibrium displacement towards the deck model as observed in the SAXS and NMR analyses. Conversely, when weakening RAR binding site using an RAR agonist or the RARI396E mutant, there is a descrease in the overall affinity of RXR/RAR for N-CoR_NID_, as a consequence of the reduction of the population of the deck complex in favour of the assymetric binding mode that eventually dissociates. Therefore, the equilibrium between the two interaction modes has relevant consequences for gene transcription, the assymetric mode facilitating coregulator swapping from corepressor to coactivator. Thus, despite the high overall affinity of the corepressor complex in the deck binding mode, the low affinity of the individual CoRNR motifs for the LBDs permits effective coregulator binding versatility upon environmental perturbations. The multivalency of coregulator proteins has important consequences in the thermodynamics of the interaction and the kinetics of the transition between repressed and active states. Multivalency is specially relevant for NR regulation as coactivators and corepressors have been described to contain several LBD interaction boxes with slightly different individual affinities. As a consequence, the number of coexisting assembly states increases dramatically, complexifying the regulation mechanism. In this context, the integrative approach applied in the present study will offer the opportunity to disentangle this complexity, structurally characterize the individual states and address the thermodynamics of NR gene transcription regulation.

## Methods

### Ligands and peptides

The ligands Am580, BMS493 and TTNPB were purchased from Tocris Bioscience. The fluorescent peptides CoRNR1 (fluorescein-RLITLADHICQIITQDFAR) and CoRNR2 (fluorescein-DPASNLGLEDIIRKALMGSFD from N-CoR were purchased from EZbiolab.

### Two-hybrid experiments

COS cells were cultured in DMEM with Glutamax and 5% (v/v) FCS and transfected using JetPei transfectant (Ozyme). After 24 h, the medium was changed to a medium containing the indicated ligands or vehicle. Cells were lysed and assayed for reporter expression 48 h after transfection. The luciferase assay system was used according to the maunfacturer’s instruction (Promega). In each case, results were normalized to coexpressed β –galactosidase. Each transfection was carried out in duplicate and repeated three to six times each.

### Constructs, expression and purification of proteins and complexes

The heterodimer RXR/RAR (mRXRα/hRARα) was prepared as previously described in (Pogenberg et al., 2005). The heterodimer RXRΔH12/RAR (mRXRαΔH12/hRARα) comprises a shorter form of RXR in which the region N227 to D444 including helix H12 is deleted. The heterodimer RXR/RARI396E (mRXRα/hRARαI396E) comprises the I396E mutation of RARα (Le Maire et al., 2010). The N-CoR_NID_ protein corresponds to the sequence from Gln2059 to Glu2325 of mouse N-CoR and was prepared as described in (Harrus et al., 2018). For the preparation of the complexes, the purified heterodimers (RXR/RAR or RXRΔH12/RAR) were mixed with a two-fold molar excess of purified N-CoR_NID_ fragment and incubated overnight at 4°C. Then, the mix was loaded on a S200 superdex gel filtration column in a buffer consisting in 50mM Tris HCl pH7.5, 150mM NaCl, 2mM DTT and fractions corresponding to the ternary complexes were pooled. The purity of the complexes was checked on a SDS-page gel.

### Steady-state fluorescence anisotropy

We performed assays using a Safire microplate reader (TECAN) with the excitation wavelength set at 470 nm and emission measured at 530 nm for fluorescein-tagged peptides. The buffer solution for assays was 20 mM Tris-HCl, pH 7.5, 150 mM NaCl, 1 mM EDTA, 5 mM DTT and 10% (v/v) glycerol. We initiated the measurements at the highest concentration of protein (20 µM) and diluted the protein sample successively two-fold with the buffer solution. For each point of the titration curve, we mixed the protein sample with 4 nM of fluorescent peptide and 40 µM of ligand (two molar equivalents). We fitted binding data using a sigmoidal dose-response model (GraphPad Prism, GraphPad Software). The reported data are the average of at least three independent experiments.

### MicroScale Thermophoresis (MST)

A fluorescent probe (Atto647N maleimide, Invitrogen) was attached to the thiol group of the sole cysteine of the N-CoR_NID_ fragment, according to the manufacturer’s instructions. This cysteine (Cys 2074) is in the CoRNR1 motif but is not involved in the interaction between RAR and N-CoR as it points to the outside in the crystal structure of the complex (Le Maire et al., 2010). Heterodimers (RXR/RAR, RXRΔH12/RAR and RXR/RARI396E) were prepared as a twofold serial dilution in MST buffer (NanoTemper Technologies GmbH) and added to an equal volume of 80nM labelled N-CoR_NID_, in MST buffer. After 10 min incubation time, the complex was filled into Monolith NT.115 Premium Coated Capillaries (NanoTemper Technologies GmbH) and thermophoresis was measured using a Monolith NT.115 Microscale Thermophoresis device (NanoTemper Technologies GmbH) at an ambient temperature of 22°C, with 5 s/30 s/ 5 s laser off/on/off times, respectively. Instrument parameters were adjusted with 40% red LED power and 20% IR-laser power. Data from three independent measurements were analyzed (NT Analysis software last version, NanoTemper Technologies GmbH) using the signal from Thermophoresis + T-Jump.

### Analytical ultracentrifugation sedimentation velocity experiments (AUC-SV)

Samples of N-CoR_NID_, RXR/RAR, N-CoR_NID_:RXR/RAR, N-CoR_NID_:RXR/RAR in the presence of BMS493, and N-CoR_NID_:RXRΔH12/RAR at about 1 g/L in 50 mM Tris pH 7.4, 150 mM NaCl, 2 mM DTT, were investigated in SV experiments. SV experiments were conducted in an XLI analytical ultracentrifuge (Beckman, Palo Alto, CA) using an ANTi-50 rotor and detection at 280 and 295 nm, at 42 000 revs. per minutes (rpm) (130 000g) and 12°C overnight, using double channel center pieces (Nanolytics, Germany) of 12 mm optical path length (loaded volume: 400µL, the reference channel being filled with the solvent) equipped with sapphire windows. For each sample, because the total absorbance at 280 nm exceeded 1.2, the set of SV profiles obtained at 295 nm was analyzed, using analysis in terms of a continuous distribution, *c*(*s*), of sedimentation coefficients, *s*, and of one non interacting species, providing independent estimated of *s* and the molar mass, *M*, of the Sedfit software (Schuck 2000), version 14.1 (freely available at: http://www.analyticalultracentrifugation.com). The related figures were made with the program Gussi (Brautigam et al., 2016) (freely available at: http://biophysics.swmed.edu/MBR/software.html). The values of *s* were corrected for experimental conditions (12°C, solvent density, *ρ*° =1.007 g/mL, and viscosity, *η*° =1.235 g/mL, estimated using the program Sednterp (freely available at: http://sednterp.unh.edu/) to values, *s*_20w_, at 20°C in pure water (*p°* =0.99823 g/mL, and viscosity, *η*° =1.002 g/mL), and interpreted, through the Svedberg equation *s = M* (1 - *p*° 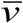) / (*N*_A_6*π η*° *R*_H_), where *M* is the molecular mass, 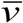 is the partial specific volume (0.720 mL/g for N-CoR_NID_, 0.743 mL/g for RXR/RAR complexes, 0.735 mL/g for N-CoR_NID_:RXR/RAR complexes, calculated using the program Sedfit), and *N*_A_ is Avogadro’s number. *R*_H_ is the hydrodynamic radius. *R*_H_ = *f/f*_min_ *R*_min_, with *f/f*_min_ the frictional ratio and *R*_min_ the radius of the anhydrous volume. The *s*20w-values were analyzed in term of *f/f*_min_ considering the the molar masses calculated from different association states. Stoichiometries with *f/f*_min_ in the range 1.25 to 1.5 were considered as acceptable, since corresponding to globular compact to reasonably anisotropic/elongated or extended shapes. The results of the non-interacting species analysis provided molar masses about 20% lower than the derived stoichiometry, which can reasonably be related to the low signal/noise level of the data.

### Mass spectrometry (RPLC-UV/MS)

Relative abundance of RAR, RXR and N-CoR_NID_ in purified complex was assessed using reverse phase liquid chromatography (Waters Alliance 2695) coupled to UV (diode array detector) and MS (Waters LCT operated in the positive ion mode) detection. Protein sample was diluted in buffer A (0.1% TFA / 0.1% FA /water) before 600 pmol of complex were injected onto reverse phase Xbridge BEH300 C4 column (3.5μm; 50 × 2.1mm; T 65°C; flow rate 0.2 ml/min). Eluting gradient was ramped from 2 to 90% buffer B (0.08% TFA / 0.1% FA / 70% ACN / 30% IprOH) in 40 min. Eluting species were detected by UV and mass spectrometry. Relative abundance of RAR, RXR and N-CoR_NID_ was calculated from protein theoretical extinction coefficient and UV peak area (λ = 280 nm) according to equation: *Abs*_280*nm*(*P*)_ = *ɛ*_280 *nm*(*P*)_ X L X [*P*] where “Abs” and “ɛ” correspond to protein (P) absorbance and extension coefficient (M^−1^ cm^−1^), respectively, at 280 nm. “L” is the light path length (cm) and [P] corresponds to protein concentration (M).

### Structural bioinformatics on N-CoR_NID_

Local secondary structural features of N-CoR_NID_ were predicted using the support vector machine-based PSIPRED3.2 server (Buchan et al., 2013). Disorder propensity of N-CoR_NID_ was assessed with IUPRED (Dosztányi et al., 2005), PrDOS (Ishida and Kinoshita, 2007), PONDR-FIT (Xue et al., 2010) and DISOPRED3 (Jones and Cozzetto, 2015). The GREMLIN software (Kamisetty et al., 2013) was fed with the N-CoR_NID_ sequence to capture evolutionary conserved sequences by searching a pre-clustered Uniprot database using the HHblits algorithm (Remmert et al., 2012). A Hidden Markov Model (HMM) multiple sequence alignment (MSA) was generated excluding hits with ≥ 75 % gaps. A final set of 61 sequences was used for detecting conservation and co-evolution signatures at residue-level. Conservation profile (Figure 2G) was extracted from the HMM MSA using Skylign to compute the probability of observing for a given position a specific residue above the background (Wheeler et al., 2014). Co-evolution analysis within the MSA was performed using BIS^2^ (Oteri et al., 2017), which bypass the statistics requirements to estimate the background noise and the relevance of the co-evolution signal. BIS^2^ is a combinatorial method that allows identifying co-evolving blocks, instead of only residues, in a small number of homologous sequences.

### Small-angle X-ray scattering

Small-angle X-ray scattering data were measured at the BM29 beam-line using the X-ray wavelength of 0.99Å and a sample-to-detector distance of ~2.9 m (Pernot et al., 2013). Several datasets were collected at multiple concentrations (Table S1 and S3) in 50mM Tris HCl pH7.5, 150mM NaCl, 2mM TCEP. Prior to data collection, the isolated proteins and the ternary complexes were supplemented with 2mM TCEP and concentrated. When necessary, ligands (BMS493 and Am580) were added to the complex in a three molar excess before concentration. Repetitive measurements allowed to detect and to correct for radiation damage and scattering patterns of the buffer solutions were recorded before and after the measurements of each protein sample. Final curves for each concentration were derived after subtracting the averaged buffer scattering from the protein sample patterns using PRIMUS. SAXS curves of different concentrations were merged to avoid interparticle interactions using standard protocols with ATSAS software (Franke et al., 2017).

The *R_g_* values in Table S1 and Table S3 were estimated by applying the Guinier approximation in the range *s <* 1.3/*R_g_* for globular proteins and *s <* 0.8/*R_g_* for free N-CoR_NID_. Pair-wise distance distribution functions, *P(r)*, were obtained by indirect Fourier Transform of the scattering intensities with GNOM (Svergun et al., 1988). We used CRYSOL (Svergun et al., 1995) to compute the theoretical SAXS profiles from conformational ensembles of free N-CoR_NID_, RXR/RAR, and N-CoR_NID_:RXR/RAR complexes. All theoretical curves were obtained with 101 points, and a maximum scattering vector of 0.5 Å^−1^ using 25 harmonics. The resulting ensemble-based CRYSOL SAXS curves, or their weighted combination, were directly compared to the corresponding experimental SAXS data with OLIGOMER (Konarev et al., 2003).

### Nuclear Magnetic Resonance (NMR)

NMR data were recorded using 140 µM U-[^15^N] N-CoR_NID_ and 600 µM U-[^13^C/^15^N] N-CoR_NID_ labeled protein in 50mM Bis Tris buffer at pH 6.7, 150mM NaCl, 1mM EDTA and 2mM TCEP with 7% (v/v) ^2^H_2_O. NMR assignment experiments were recorded at 293 K on a 950 MHz Bruker Avance III spectrometer equipped with a cryogenic triple-resonance probe (IR-RMN Gif/Yvette). ^1^H chemical shifts were referenced directly, and ^15^N chemical shifts indirectly (Markley et al., 1998), to added 2,2-dimethyl-2- silapentane-5-sulfonate (DSS, methyl ^1^H signal at 0.00 ppm). A set of 3D Best-TROSY HNCO, HN(CA)CO, HN(CO)CA, HNCA, iHNCA, HN(CO)CACB, HNCACB, iHNCACB (Lescop et al., 2007), and H-N-N experiments HN(CA)NH, HN(COCA)NH. NMR interaction and PRE experiments (Barrett et al., 2013) were recorded at 293 K on a 700 MHz Bruker Avance III spectrometer equipped with a cryogenic triple-resonance probe. Spectra were processed with nmrpipe and analyzed using nmrview (Johnson and Blevins, 1994). NMR ^1^H-^15^N HSQC’s have been recorded for ^15^N labelled N-CoR_NID_, and for ^15^N-N-CoR_NID_:RXR/RAR (1:1), ^15^N-N-CoR_NID:_RXRΔH12/RAR (1:1), ^15^N-N-CoR_NID_: RXR/RARI396E (1:1.2). ^1^H-^15^N HSQCs were also measured for equimolar complexes of ^15^N-N-CoR_NID_:RXR/RAR in the presence of 1.2 molar excess of RAR agonist (Am580) or RAR inverse agonist (BMS493). For PRE measurements, ^1^H-^15^N HSQC spectra of diamagnetic and paramagnetic samples of ^15^N-N-CoR_NID_:RXR/RAR (1:1.3) were acquired with a recycling delay between scans of 2 s to ensure that magnetization recovery level is identical for both states and using the same concentration and number of scans for both samples. The paramagnetic contribution to the relaxation rate was determined as the ratio of peak intensities in the paramagnetic and diamagnetic states.

### Site-directed spin-labeling for paramagnetic studies

In addition to the wild-type cysteine residue C2074, two additional single-cysteine variants of N-CoR_NID_ (S2213C and S2288C) were engineered by site-directed mutagenesis after substitution of the native cysteine by a serine (C2074S), using QuikChange Lightning Site-Directed mutagenesis kit. All constructs were verified by DNA sequencing. In order to conjugate the paramagnetic tag, a 10-fold excess of 3-(2-Iodoacetamido)-2,2,5,5-tetramethyl-1-pyrrolidinyloxy (IA-PROXYL) spin label was immediately added to reducing agent-free samples, just after elution from PD-10 desalting columns, and left reacting for 3 h in the dark at room temperature. Excess of unreacted tag was removed by passing the reaction mixture twice through a PD-10 column, and exchanged to 50mM Bis Tris buffer at pH 6.7, 150mM NaCl, 1mM EDTA. Reference diamagnetic samples were obtained after adding 5-fold of fresh ascorbic acid to the same sample used to acquire the paramagnetic spectra.

### Structural modeling of RXR/RAR heterodimer

Multiple structural all-atoms models of the RXR/RAR were built in order to describe SAXS curves of the heterodimer in different experimental conditions. The fully disordered H12 scenario was built using the 1DKF structure as a template and by adding the disordered N- and C-termini (including the H12) extensions as well as the connecting linker between helices H1 and H3. Previous crystallographic structures were used to model the other scenarios where one (RAR) or both (RAR and RXR) H12 regions were placed folded and bound to the rest of the LDB in either an agonist or antagonist conformation. Concretely, the structure 1DKF (Bourguet et al., 2000) was used as a template for the antagonist conformation, and the structures 1XDK (Pogenberg et al., 2005) and 3KMR (Le Maire et al., 2010) were used as templates for the agonist conformation (Fig. S1 and Table S2). Ensembles of disordered fragments were built using *Flexible-Meccano* (FM) (Bernado et al., 2005; Ozenne et al., 2012) and attached to the X-ray templates using *in-house* scripts (Cordeiro et al., 2017b). Briefly, for each disordered segment built, side-chains were added using SCCOMP (Eyal et al., 2004) and then pre-processed with Rosetta 3.5 *fixbb* module (Kuhlman et al., 2003) to alleviate steric clashes. Each structure was then refined in explicitly solvent using Gromacs 5.0.2 (Hess et al., 2008). All ensembles of RXR/RAR heterodimer comprised 5,000 conformers.

### Structural ensembles of free N-CoR_NID_

A random coil ensemble model of N-CoR_NID_ containing 10,000 conformations was calculated also employing FM (Bernado et al., 2005; Ozenne et al., 2012) followed by side-chains modelling and refinement as described above. A second structurally biased ensemble derived from the evolutionary conservation and a co-evolutionary analysis was computed in the following way. A starting random coil ensemble of 200,000 conformations of N-CoR_NID_ was filtered selecting those structures presenting at least two contacts (≤ 15 Å) between residues in distal blocks of conserved (with Bits ≥ 1.6), structured (average disorder propensity below 0.5), and/or co-evolving residues (Fig. 2 and Fig. 3) within the C-terminal region. The filtered conformations (≈ 35,000) resulted in a compact ensemble that did not describe the SAXS data. We further refined it by performing 200 independent Ensemble Optimization Method (EOM) runs (Bernado et al., 2007), and those conformations that were not selected in any of them were discarded. The SAXS-refined ensembles was more extended than that obtained by distance restraints. The final ensemble of 10,000 conformations was use to interpret the PRE data. Briefly, for each conformer, the theoretical intramolecular ^1^H-^15^N-PRE-rates were computed using the Solomon-Bloembergen approximation (Iwahara et al., 2004; Salmon et al., 2010). Dynamics of the paramagnetic PROXYL moiety was accounted for by using multiple conformational states derived from a 100 ns Molecular Dynamics (MD) simulation of a Gly-Gly-**Cys**-Gly-Gly peptide in water at 298K, where the PROXYL moiety was attached to the central cysteine residue (Beck et al., 2008; Polyhach et al., 2011). From this library, the native and engineered single-cysteine residues were *in-silico* labelled with multiple (60) sterically allowed spin-label dispositions. This ensemble representation enabled the estimation of the order parameters that account for the motion of the dipolar proton-electron interaction vector (Iwahara et al., 2004; Salmon et al., 2010). The correlation time of N-CoR_NID_ was estimated to be 5.79 ns using HYCUD (Parigi et al., 2014). With this strategy, the PRE-rates for C2074, S2213C and S2288C were independently calculated for each N-CoR_NID_ conformer, averaged over the complete ensembles of the three N-CoR_NID_ forms, and compared with the experimental ones without any optimization process (Salmon et al., 2010).

### Molecular Dynamics Simulation

Simulations were based on the one-bead-per-residue coarse-grained model proposed by (Smith et al., 2014) for intrinsically disordered proteins. N-CoR_NID_ fragments with high α-helix propensity (2065-2088, 2214-2234, 2269-2291) were restrained to helical conformation *via* an elastic network with a force constant of 500 kJ.mol^−1^ .nm^−2^ whereas the original bonded potential for disordered chains was used for the remaining regions. Electrostatic interactions between charged residues were represented with a Debye Huckel energy functions (λ = 0.9 nm and ɛES = 1.485). Excluded volume and attractive hydrophobic interactions were modeled combining a purely repulsive Weeks-Chandler-Andersen (WCA) potential with the attractive part of a Lennard Jones potential (σ = 0.58 nm). The parameter αCG, which determines the strength of the hydrophobic interactions, was specifically tuned for N-CoR_NID_ system by minimizing the χ^2^ between experimental and simulated SAXS curves. The latter were calculated from coarse-grained trajectories using CRYSOL (Svergun et al., 1995) upon reconstruction of the full atomistic structure via BBQ (Gront et al., 2007) and SCWRL4 (Krivov et al., 2009). Notably, this procedure revealed that an optimal agreement with SAXS data (χ^2^ = 0.87) was obtained for αCG = 0.5, which is remarkably similar to the result obtained in the original publication for a set of diverse disordered proteins by comparison with FRET data. All simulations were performed with the GROMACS 5.4 molecular dynamics package (Abraham et al., 2015). An extended configuration was generated and used as initial structure for the simulation inside a 78733 nm^3^ cubic box with periodic boundary conditions. The equations of motion were integrated every 20 fs during stochastic dynamics with a damping coefficient set to 25 amu.ps^−1^. A plain cutoff of 5.0 nm was used for both Lennard-Jones and electrostatic interactions. All of the simulations were performed in the NVT ensemble and the temperature was kept constant to a reference value of 300 K. During the production, five replicas of 1 µs each were run for a total simulation length of 5 µs.

### Structural Modelling of N-CoR_NID_:RXR/RAR complexes

The structure of full-length N-CoR_NID_ bound to RXR/RAR was modeled exploiting the existing crystal structure of the RAR/NCoR CoRNR1 complex (3KMZ) (Le Maire et al., 2010) and that of the RXR in complex with SMRT CoRNR2 complex (3R29) (Zhang et al., 1999), which has a 75% of sequence identity with NCOR CoRNR2. Using S-CoRNR2 as a template we created a homology model for bound CoRNR2. In the complexes, the conformations of bound CoRNR1 and/or CoRNR2 were maintained as in the crystal structures. For singly bound complexes (asymmetric), with N-CoR_NID_ bound to RAR or RXR through CoRNR1 or CoRNR2 respectively, disordered statistical coil N- and C-terminal extensions were built onto the structured peptide employing the above-described work-flow to model ensembles of flexible fragments (Cordeiro et al., 2017a). The doubly bound complex was created in two steps. Firstly, we use FM to generate a structural ensemble of 100,000 singly bound complexes with N-CoR_NID_ bound to RXR/RAR through CoRNR1, but with CoRNR2 structured as in the crystal. Secondly, this starting ensemble was refined by selecting those conformers with CoRNR2 in the vicinity of RXR binding site (i.e. ≤ 6.5), followed by a docking process driven by the surface contacts observed in the RXR/SMRT CoRNR2 complex (PDB:3R29), which were integrated as distance restraints in the HADDOCK docking approach (Dominguez et al., 2003). Structures were then ranked using the energy-based HADDOCK scoring function and a term quantifying the RMSD of the binding interface with the RXR/SMRT CoRNR2 complex. In the apo-form and in the presence of BMS493, RAR/RXR was modelled with disordered H12s. For RAR/RXRI396E mutant and in the presence of Am580, RAR H12 was placed in the agonist position, and RXR H12 was removed in the complex between N-COR_NID_ and RXRΔH12/RAR. All ensembles of N-CoR_NID_:RXR/RAR complexes contained 2,000 conformers.

## Acknowledgments

The CBS is a member of the France-BioImaging (FBI) and the French Infrastructure for Integrated Structural Biology (FRISBI), 2 national infrastructures supported by the French National Research Agency (ANR-10-INBS-04-01 and ANR-10-INBS-05, respectively). We acknowledge the platforms of the Grenoble Instruct center (ISBG; UMS 3518 CNRS-CEA-UGA-EMBL) supported by the French Infrastructure for Integrated Structural Biology Initiative FRISBI (ANR-10-INSB-05-02) and by the Grenoble Alliance for Integrated Structural Cell Biology GRAL (ANR-10-LABX-49-01) within the Grenoble Partnership for Structural Biology (PSB). We acknowledge the use of BioSAXS BM29 beamline at ESRF-Grenoble. Financial support from the TGIR-RMN-THC Fr3050 CNRS for conducting the research is gratefully acknowledged. We acknowledge the financial support from the ANR GPCteR (ANR-17-CE11-0022 to NS). This work has benefited from support by the Labex EpiGenMed, an «Investissements d’avenir» program, reference ANR-10-LABX-12-01. We acknowledge the Laboratory of Spectroscopy and Calorimetry (LEC) at Brazilian Biosciences National Laboratory (LNBio), CNPEM, Campinas, Brazil for their support with the use of equipment (Monolith NT.115 Microscale Thermophoresis device (NanoTemper Technologies GmbH). We acknowledge the financial support from the CNPq Programa Ciencia Sem Fronteiras (BJT 300143/2015-0 to ALM) and from the CNPq Programa Universal (420416/2016-1 to ALM). We acknowledge financially support from FEDER-COMPETE2020 and FCT (Project LISBOA-01-0145-FEDER-007660 to TNC).

